# SCellBOW: AI-Driven Tumor Risk Stratification from Single-Cell Transcriptomics Using Phenotype Algebra

**DOI:** 10.1101/2022.12.28.522060

**Authors:** Namrata Bhattacharya, Anja Rockstroh, Sanket Suhas Deshpande, Sam Koshy Thomas, Anunay Yadav, Chitrita Goswami, Smriti Chawla, Pierre Solomon, Cynthia Fourgeux, Gaurav Ahuja, Brett Hollier, Himanshu Kumar, Antoine Roquilly, Jeremie Poschmann, Melanie Lehman, Colleen C. Nelson, Debarka Sengupta

**Affiliations:** Australian Prostate Cancer Research Centre-Queensland, Faculty of Health, School of Biomedical Sciences, Centre for Genomics and Personalised Health, Queensland University of Technology, Brisbane, Queensland-4000, Australia; Center for Computational Biomedicine, Harvard Medical School, Boston, MA-02115, USA; Laboratory of Immunology and Infectious Disease Biology, Department of Biological Sciences, Indian Institute of Science Education and Research (IISER), Bhopal, India; Department of Computational Biology, Indraprastha Institute of Information Technology-Delhi (IIIT-Delhi), Okhla, Phase III, New Delhi-110020, India; Department of Computer Science and Engineering, Indraprastha Institute of Information Technology-Delhi (IIIT-Delhi), Okhla, Phase III, New Delhi-110020, India; Centre for Artificial Intelligence, Indraprastha Institute of Information Technology-Delhi (IIIT-Delhi), Okhla, Phase III, New Delhi-110020, India; Nantes Université, CHU Nantes, INSERM, Center for Research in Transplantation and Translational Immunology, UMR, 1064, Nantes, France; Translational Research Institute, Princess Alexandra Hospital, Woolloongabba, Queensland-4102, Australia; Vancouver Prostate Centre, Department of Urologic Sciences, University of British Columbia, Vancouver, Canada; School of Mathematical Sciences, The University of Adelaide, North Terrace, Adelaide, SA-5005, Australia

**Keywords:** Intra-tumor heterogeneity, single-cell RNA-seq, risk stratification, transfer learning, clustering, *phenotype algebra*, prostate cancer

## Abstract

Single-cell RNA-sequencing (scRNA-seq) coupled with robust computational analysis facilitates the characterization of phenotypic heterogeneity within tumors. Current scRNA-seq analysis pipelines are capable of identifying a myriad of malignant and non-malignant cell subtypes from single-cell profiling of tumors. However, given the extent of intra-tumoral heterogeneity, it is challenging to assess the risk associated with individual cell subpopulations, primarily due to the complexity of the cancer phenotype space and the lack of clinical annotations associated with tumor scRNA-seq studies. To this end, we introduce SCellBOW, a scRNA-seq analysis framework inspired by document embedding techniques from the domain of Natural Language Processing (NLP). SCellBOW is a novel computational approach that facilitates effective identification and high-quality visualization of single-cell subpopulations. We compared SCellBOW with existing best practice methods for its ability to precisely represent phenotypically divergent cell types across multiple scRNA-seq datasets, including our in-house generated human splenocyte and matched peripheral blood mononuclear cell (PBMC) dataset. For tumor cells, SCellBOW estimates the relative risk associated with each cluster and stratifies them based on their aggressiveness. This is achieved by simulating how the presence or absence of a specific cell subpopulation influences disease prognosis. Using SCellBOW, we identified a hitherto unknown and pervasive AR−/NE_low_ (androgen-receptor-negative, neuroendocrine-low) malignant subpopulation in metastatic prostate cancer with conspicuously high aggressiveness. Overall, the risk-stratification capabilities of SCellBOW hold promise for formulating tailored therapeutic interventions by identifying clinically relevant tumor subpopulations and their impact on prognosis.

## INTRODUCTION

Intra- and inter-tumoral heterogeneity are pervasive in cancer and manifest as a constellation of molecular alterations in tumor tissues. The late-stage clonal proliferation, partial selective sweeps, and spatial segregation within the tumor mass collectively orchestrate lineage plasticity and metastasis^1^. In collaboration with non-malignant cell types in the tumor stroma, malignant cells with distinct genetic and phenotypic properties create complex and dynamic ecosystems, rendering the tumors recalcitrant to therapies^2^. Thus, the phenotypic characterization of malignant cell subpopulations is critical to understanding the underlying mechanisms of resistive behavior. The widespread adoption of single-cell RNA-sequencing (scRNA-seq) has enabled the profiling of individual cells, thereby obtaining a high-resolution snapshot of their unique molecular landscapes^3,4^. A precise understanding of cell-to-cell functional variability captured by scRNA-seq profiles is crucial in this context. These molecular profiles assist in robust deconvolution of the oncogenic processes instigated by various selection pressures exerted by anticancer agents. They also facilitate understanding the cross-talks between malignant and non-malignant cell types within the tumor microenvironment^5^. To effectively analyze tumor scRNA-seq data, various specialized techniques have been developed. These techniques assist in proactive investigation of complex and elusive cell populations^6,7^, regulatory gene interactions^8^, neoplastic cell lineage trajectories^9^, and expression-based inference of copy number variations^10^. While these computational techniques have been successful in gaining novel biological insights, their adoption in clinical settings is still elusive.

Over the past years, leading consortia such as The Cancer Genome Atlas (TCGA)^11^ and other large-scale independent studies have established reproducible molecular subtypes of cancers with divergent prognoses. For instance, metastatic prostate cancer has typically been categorized based on the androgen receptor (AR) activity or neuroendocrine (NE) program: the less aggressive AR+/NE– (AR prostate cancer, ARPC) and the highly aggressive AR–/NE+ (NE prostate cancer, NEPC)^12^. More recent studies have identified additional phenotypes, such as the AR–/NE– (double-negative prostate cancer, DNPC) and AR+/NE+ (amphicrine prostate cancer, AMPC)^13,14^. These findings underscore the importance of considering tumor heterogeneity while dictating the differential treatment regimes to improve patient outcomes^15^. These studies are predominantly based on bulk omics assays, which precludes the detectability of fine-grained molecular subtypes of clinical relevance. To address this, there is an urgent need to develop novel analytical approaches that are capable of exploiting single-cell omics profiles for risk attribution to malignant cell subtypes.

A growing number of NLP-based methods have recently been gaining popularity for their ability to predict patients’ survival based on their clinical records and for facilitating efficient analyses of high-dimensional scRNA-seq data^16–19^. For instance, Nunez *et al.*^16^ employ language models to predict the survival outcomes of breast cancer patients based on their initial oncologist consultation documents. Similarly, in the domain of scRNA-seq analysis, scETM^18^ uses topic models to infer biologically relevant cellular states from single-cell gene expression data. Recently, scBERT^19^, an adaptation of attention-based transformer architecture, has been developed for cell type annotation in single-cell. Motivated by the objective to efficiently associate survival risk directly with cellular subpopulations extracted from scRNA-seq data, we introduce SCellBOW (single-cell bag-of-words), a Doc2vec^20^ inspired transfer learning framework for single-cell representation learning, clustering, visualization, and relative risk stratification of cell types within a tumor microenvironment. SCellBOW intuitively treats cells as documents and genes as words. SCellBOW learned latent representations capture the semantic meanings of cells based on their gene expression levels. Due to this, cell type or condition-specific expression patterns get adequately captured in cell embeddings. We demonstrate the utility of SCellBOW in clustering cells based on their semantic similarities across multiple scRNA-seq datasets, spanning from normal prostate to pancreas and peripheral blood mononuclear cells (PBMC). As an extended validation, we apply SCellBOW to an in-house scRNA-seq dataset comprising human PBMC and matched splenocytes. We observed that SCellBOW outperforms existing best-practice single-cell clustering methods in its ability to precisely represent phenotypically divergent cell types.

Beyond robust identification of cellular clusters, the latent representation of single cells provided by SCellBOW captures the ‘semantics’ associated with cellular phenotypes. In line with *word algebra* supported by NLP models, which explores word analogies based on semantic closeness (e.g., adding a word vector associated *royal* to *man* brings it closer to *king*), we conjectured that cellular embedding can reveal biologically intuitive outcomes through algebraic operations. Thus, we aimed to replicate this feature in the single-cell phenotype space to introduce *phenotype algebra* and apply the same to attribute prognostic value to cancer cell subpopulations identified from scRNA-seq studies. With the help of *phenotype algebra,* it is possible to simulate the exclusion of the phenotype associated with a specific subpopulation within the tumor microenvironment and relate the same to the disease outcome. Selectively removing a specific subtype from the whole tumor allows us to categorize the aggressiveness of individual subtypes under negative selection pressure while preserving the impact of the residual tumor. This information can ultimately aid therapeutic decision-making and improve patient outcomes. As a proof of concept, we validate the *phenotype algebra* module of SCellBOW for risk-stratification of malignant and non-malignant cell subtypes across three independent cancer types: glioblastoma multiforme, breast cancer, and metastatic prostate cancer.

Finally, we demonstrate the utility of SCellBOW to uncover and define novel subpopulations in a cancer type based on biological features relating to disease aggressiveness and survival probability. Metastatic prostate cancer encompasses a range of malignant cell subpopulations, and characterizing novel or more fine-grained subtypes with clinical implications on patient prognosis is an active field of research^21,22^. To explore this, we applied our *phenotype algebra* on the SCellBOW clusters as opposed to predefined subtypes. Through our analysis using SCellBOW, we identify a dedifferentiated metastatic prostate cancer subpopulation. This subpopulation is nearly ubiquitous across numerous metastatic prostate cancer samples and demonstrates greater aggressiveness compared to any known molecular subtypes. To the best of our knowledge, SCellBOW pioneered the application of a NLP-based model to attribute survival risk to malignant and non-malignant cell subpopulations from patient tumors. SCellBOW holds promise for tailored therapeutic interventions by identifying clinically relevant subpopulations and their impact on prognosis.

## RESULTS

### Overview of SCellBOW

Correlating genomic readouts of tumors with clinical parameters has helped us associate molecular signatures with disease prognosis^23^. Given the prominence of phenotypic heterogeneity in tumors, it is important to understand the connection between molecular signatures of cellular populations and disease aggressiveness. Existing methods map tumor samples to a handful of well-characterized molecular subtypes with known survival patterns primarily obtained from bulk RNA-seq studies. However, bulk RNA-seq measures the average level of gene expression distributed across millions of cells in a tissue sample, thereby obscuring the intra-tumoral heterogeneity^24^. This limitation prompted us to turn to tumor scRNA-seq. Tumor scRNA-seq studies have successfully revealed the extent of gene expression variance across individual malignant cells, contributing to an in-depth understanding of the mechanisms driving cancer progression^25^. However, to date, most studies use marker genes identified from bulk RNA-seq studies to characterize malignant cell clusters identified from scRNA-seq data. This approach is inadequate since every tumor is unique when looked through the lenses of single-cell expression profiles^26^. Bulk markers alone cannot adequately capture the intricate heterogeneity present within tumors, as different tumors may exhibit distinct phenotypes with varying degrees of aggressiveness at the single-cell level. While existing methods effectively reveal the subpopulations, they are insufficient in associating malignant risk with specific cellular subpopulations identified from scRNA-seq data. The proposed approach, SCellBOW, can effectively capture the heterogeneity and risk associated with each phenotype, enabling the identification and assessment of malignant cell subtypes in tumor microenvironment directly from scRNA-seq gene expression profiles, thereby eliminating the need for marker genes (**Fig. 1**). Below, we highlight key constructions and benefits of this new approach.

**Fig. 1.**
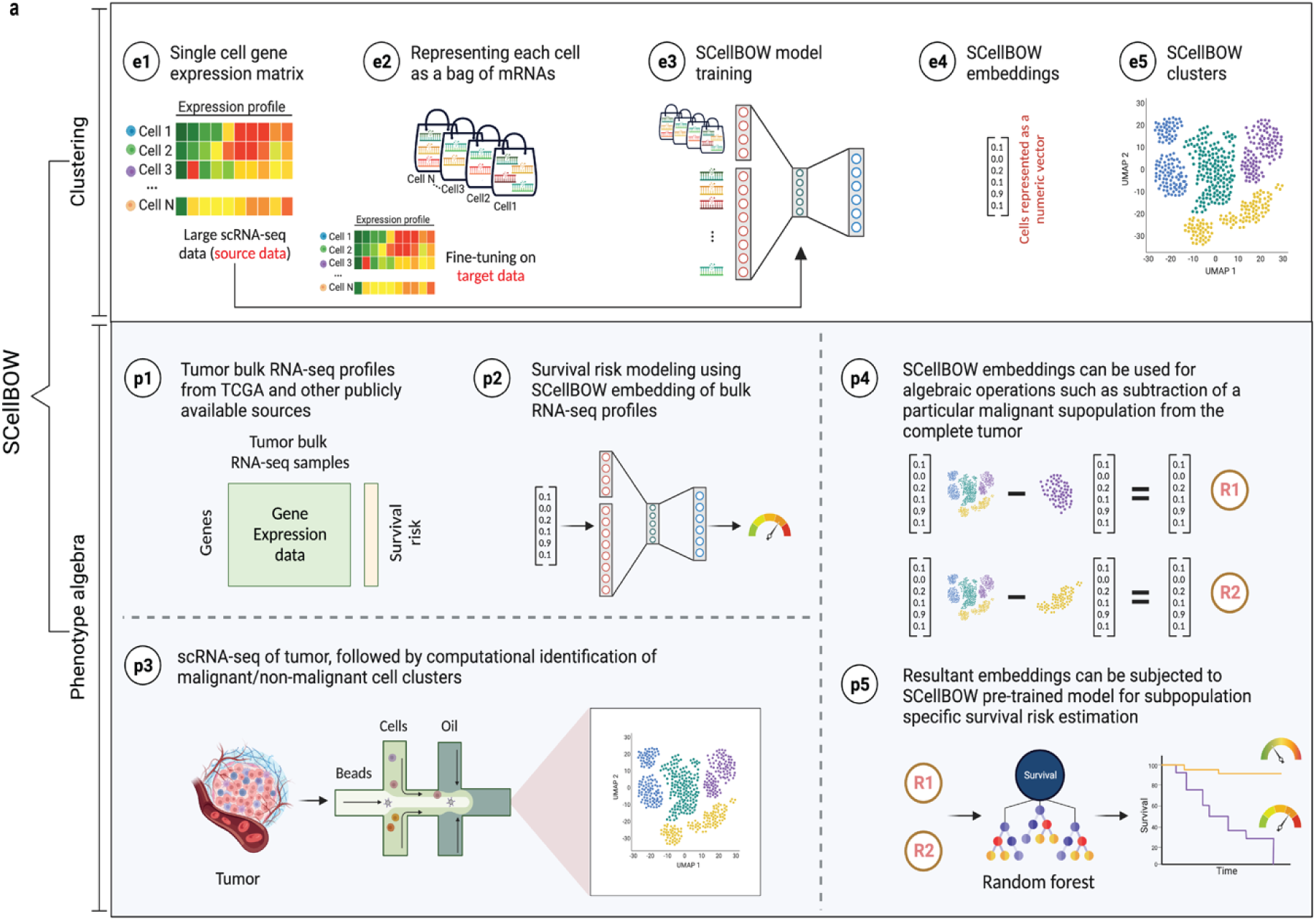
SCellBOW workflow. **a,** Schematic overview of SCellBOW workflow for identifying cellular clusters and assessing the aggressiveness of the predicted clusters. For SCellBOW clustering, firstly, a corpus was created from the gene expression matrix, where cells were analogous to documents and genes to words. Next, the pre-trained model was retrained with the vocabulary of the target dataset. Then, clustering was performed on embeddings generated from the neural network. For SCellBOW *phenotype algebra*, vectors were created for reference (*total tumor*) and queries. Then, the query vector was subtracted from the reference vector to calculate the predicted risk score using a bootstrapped random survival forest. Finally, survival probability was evaluated, and phenotypes were stratified by the median predicted risk score.

SCellBOW adapts the popular document-embedding model Doc2vec for single-cell latent representation learning, which can be used for downstream analysis. SCellBOW represents cells as documents and gene names as words. Gene expression levels are encoded by ‘word frequencies’, i.e., variation in gene expression values is captured by introducing gene names in proportion to their intensities in cells. SCellBOW encodes each cell into low-dimensional embeddings. SCellBOW extracts the gene expression patterns of individual cells from a relatively large unlabeled source scRNA-seq dataset by pre-training a shallow source network. The neuronal weights estimated during pre-training are transferred to a relatively smaller unlabeled scRNA-seq dataset, where they are refined based on the gene expression patterns in the target dataset. SCellBOW leverages the variability in gene expression values to subsequently cluster cells according to their semantic similarity.

Building upon SCellBOW’s capacity to preserve the semantic meaning of individual cells, we attempted to establish meaningful associations among cellular phenotypes. SCellBOW offers a remarkable feature for executing algebraic operations such as ‘+’ and ‘–’ on single cells in the latent space while preserving the biological meanings. This feature catalyzes the simulation of the residual phenotype of tumors, following positive and negative selection of specific cell subtype in a tumor. This could potentially be used to identify the contribution of that phenotype to patient survival. For example, the ‘–’ operation can be used to predict the likelihood of survival by eliminating the impact of a specific aggressive cellular phenotype from the whole tumor. We empirically show the retention of such semantic relationships in the context of cellular phenotypes of cancer cells using *phenotype algebra*. SCellBOW uses algebraic operations to compare and analyze the importance of cellular phenotypes working independently or in combination toward the aggressiveness of the tumor in a way that is not possible using traditional methods. *Phenotype algebra* can be performed either on the pre-defined cancer subtypes or SCellBOW clusters. Drawing from the overall performance of SCellBOW in accurately clustering and ranking cellular phenotypes by aggressiveness, we further analyzed multi-patient scRNA-seq data of metastatic prostate cancer and characterized an unknown, de-differentiated AR−/NE_low_ subpopulation of malignant cells.

### SCellBOW robustly dissects tissue heterogeneity

Malignant cells are far more heterogeneous compared to associated normal cells. Clustering is often the first step toward recognizing cellular lineages in a tumor sample. Clustering single cells based on their molecular profiles can potentially identify rare cell populations with distinct phenotypes and clinical outcomes. To evaluate the strength of the clustering ability of SCellBOW, we benchmarked our method against five existing scRNA-seq clustering methods^27–31^ (**Supplementary Table 1**). Among these packages, Seurat^27^ and Scanpy^28^ are the most popular, and both employ graph-based clustering techniques. DESC^29^ is a deep neural network-based scRNA-seq clustering package, whereas ItClust^30^ and scETM^31^ are transfer learning methods. All the packages are resolution-dependent except for ItClust. ItClust automatically selects the resolution with the highest silhouette score. We used a number of scRNA-seq datasets to evaluate the methods cited above (**Supplementary Table 2**). For the objective evaluation of performance, we used an adjusted Rand index (ARI) and normalized mutual information (NMI) to compare clusters with known cell annotations (**Supplementary Table 3)**. For all methods, except for ItClust, we computed overall ARI and NMI for different resolution values ranging between 0.2 to 2.0. We computed the cell type silhouette index (SI) based on known annotations to measure the signal-to-noise ratio of low-dimensional single-cell embeddings.

We constructed three use cases leveraging publicly available scRNA-seq datasets. Each instance constitutes a pair of single-cell expression datasets of which the source data is used for self-supervised model training and the target data for model fine-tuning and analysis of the clustering outcomes. In all cases, the target data has associated cell type annotations derived from fluorescence-activated cell sorting (FACS) enriched pure cell subpopulations. The first use case consists of non-cancerous cells from prostate cancer patients^32^ (120,300 cells) as the source data and cells from healthy prostate tissues^33^ (28,606 cells) as the target data. This use case was designed to assess the resiliency of SCellBOW to the presence of disease covariates in a large scRNA-seq dataset. The second use case comprises a large PBMC dataset^34^ (68,579 cells) as the source data, whereas the target data was sourced from a relatively small FACS-annotated PBMC dataset (2,700 cells) from the same study. The third use case, as source data, comprises pancreatic cells from three independent studies processed with different single-cell profiling technologies (inDrop^35^, CEL-Seq2^36^, SMARTer^37^). The target data used in this case was from an independent study processed with a different technology, Smart-Seq2^38^ (**Fig. 2, Supplementary Fig. 1**). For the normal prostate, PBMC, and pancreas datasets, SCellBOW produced the highest ARI scores across most notches of the resolution spectrum (0.2 to 2.0) (**Fig. 2g-i**). We observed a similar trend in the case of NMI (**Fig. 2j**). SCellBOW exhibited the highest NMI compared to other methods for the normal prostate, PBMC, and pancreas datasets. For further deterministic evaluation of the different methods across the datasets, we set 1.0 as the default resolution for calculating cell type SI (**Fig 2k**). In the PBMC and pancreas datasets, SCellBOW yielded the highest cell type SI for both datasets. SCellBOW and Seurat were comparable in performance for the pancreas dataset, outperforming other methods. We observed poor performance by DESC and ItClust across all the datasets in terms of cell type SI. In terms of cell type SI, Seurat and scETM, showed improved results in the normal prostate dataset. However, SCellBOW outperformed both Seurat and scETM in terms of overall ARI and NMI for the same dataset, indicating higher clustering accuracy. To further evaluate the cluster quality in the normal prostate dataset, we compared the known cell types to the predicted clusters. Known cell types such as basal epithelial, luminal epithelial, and smooth muscle cells were grouped into homogeneous clusters by SCellBOW (**Supplementary Fig. 2**). We observed that the majority of fibroblasts and endothelial cells were mapped by SCellBOW to single clusters, unlike Seurat and scETM. SCellBOW retained hillock cells in close proximity to both basal epithelial and club cells, in contrast to Seurat, which only includes basal epithelial cells. We compared SCellBOW with two additional methods published recently – scBERT^19^ and scPhere^39^. Among these, scBERT is a transfer learning-based method built upon BERT^40^. scPhere is a deep generative model designed for embedding single cells into low-dimensional hyperspherical or hyperbolic spaces. While scBERT offers competitive performance, scPhere consistently falls behind the other methods (**Supplementary Fig. 3 and Supplementary Note 1**).

**Fig. 2.**
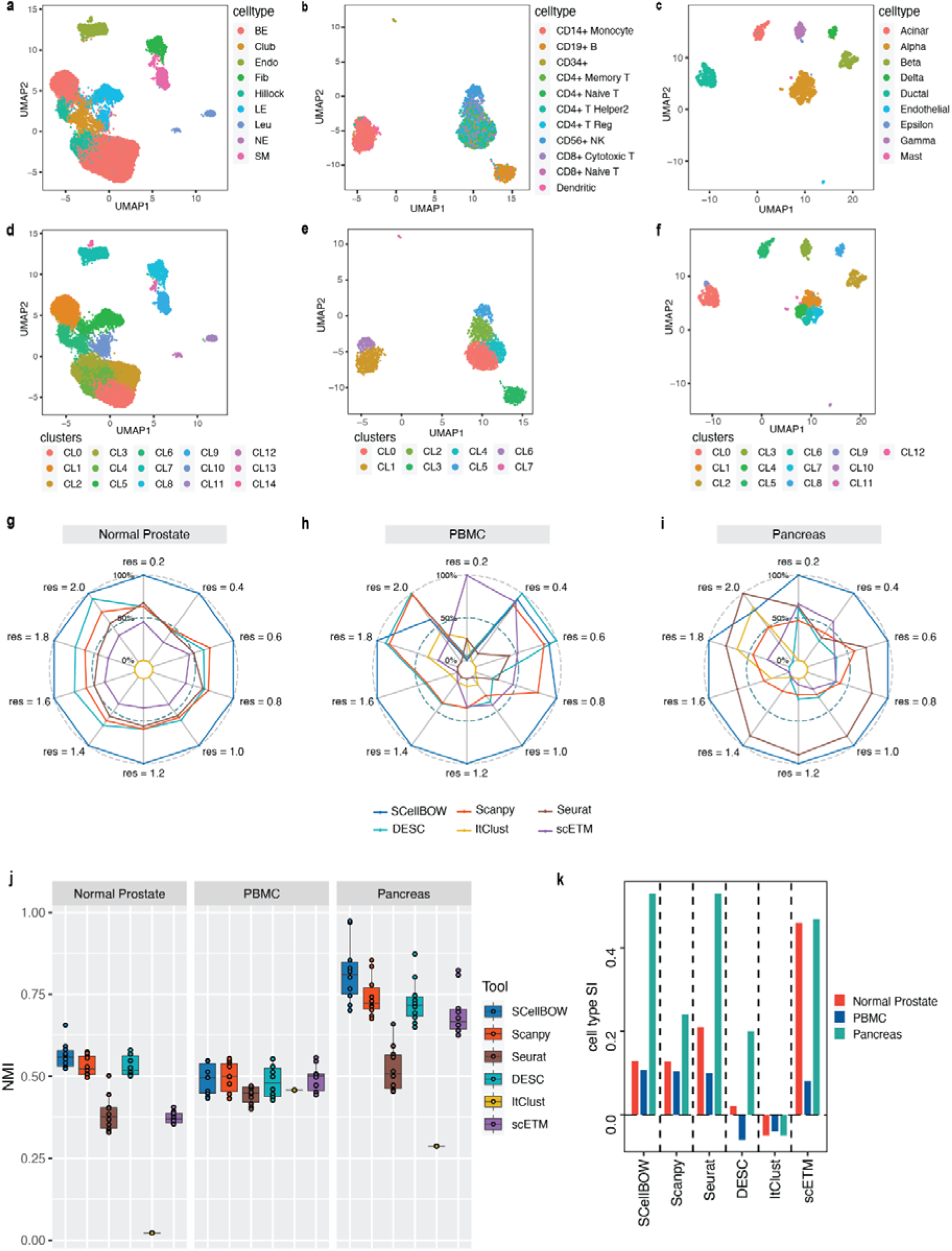
Evaluation of single-cell representations using SCellBOW. **a-c,** UMAP plots for the normal prostate (a), PBMC (b), and pancreas (c) datasets. The coordinates are colored by cell types. **d-f,** UMAP plots for normal prostate (d), PBMC (e), and pancreas (f) datasets, where the coordinates are colored by SCellBOW clusters. CL is used as an abbreviation for cluster. **g-i,** Radial plot for the percentage of contribution of different methods towards ARI for various resolutions ranging from 0.2 to 2.0. ItClust is a resolution-independent method; thus, the ARI is kept constant across all the resolutions. **j,** Box plot for the NMI of different methods across different resolutions ranging from 0.2 to 2.0 in steps of 0.2. **k,** Bar plot for the cell type silhouette index (SI) for different methods. The default resolution was set to 1.0.

Our analyses of the public datasets confirm the robustness of SCellBOW compared to the prominent single-cell analysis methods, including the prominent transfer learning methods. To this end, we applied SCellBOW to investigate a more challenging task of analyzing an in-house scRNA-seq data comprising splenocytes and matched PBMCs from two healthy and two brain-dead donors (**Fig 3a**). Given multiple covariates, such as the origin of the cells and the physiological states of the donors, analysis of this scRNA-seq data presents a challenging use case (**Fig 3b-c**). We used the established high-throughput scRNA-seq platform CITE-seq^41^ to pool eight samples into a single experiment. After post-sequencing quality control, we were left with 4,819 cells. We annotated the cells using Azimuth^42^, with occasional manual interventions (**Fig 3d, Supplementary Fig. 4d-f, Supplementary Note 2**). We quantitatively evaluated SCellBOW and the rest of the benchmarking methods by measuring ARI, NMI, and cell type SI (**Fig. 3e, Supplementary Fig. 3d-f)**. While most methods did reasonably well, SCellBOW offered an edge. We observed the best results in SCellBOW in terms of ARI, NMI, and cell type SI compared to other methods (**Supplementary Table 3**). SCellBOW yielded clusters largely coherent with the independently done cell type annotation using Azimuth (**Fig 3, Supplementary Fig. 5**). Further, most clusters harbor cells from all the donors, indicating that the sample pooling strategy was effective in reducing batch effects. B cells, T cells, and NK cells map to SCellBOW clusters where the respective cell types are the majority. Most CD4+ T cells map to CL0 and CL9 (here, CL is used as an abbreviation for cluster) (**Fig 3f**). CL0 is shared between CD8+ T cells, CD4+ T cells, and Treg cells, which originate from the same lineage. CL4 majorly consists of NK cells with a small fraction of CD8+ T cells, which is not unduly deviant from the PBMC lineage tree. As a control, we performed similar analyses of the Scanpy clusters (**Fig 3g**). While Scanpy performed reasonably well, misalignments could be spotted. For example, CD4+ and CD8+ T cells were split across many clusters with mixed cell type mappings. SCellBOW maps CD14 monocytes to a single cluster, whereas Scanpy distributes CD14 monocytes across two clusters (CL1 and CL8), wherein CL8 is equally shared with conventional Dendritic Cells (cDC). ‘Eryth’ annotated cells are indignantly mapped to different clusters by both SCellBOW and Scanpy. This could be due to the Azimuth’s reliance on high mitochondrial gene expression levels for annotating erythroid cells^42^ **(Supplementary Note 3)**. However, SCellBOW CL6 is dominated by Erythroid cells with marginal interference from other cell types. In the case of Scanpy, no such cluster could be detected. Discerning phenotypic heterogeneity from the expression profiles of seemingly similar cells is a challenging task. The performance of SCellBOW, with near-ground-truth cell type annotation, in such a scenario confirmed its ability to adequately decipher the underlying cellular heterogeneity and provide robust cell type clustering.

**Figure 3.**
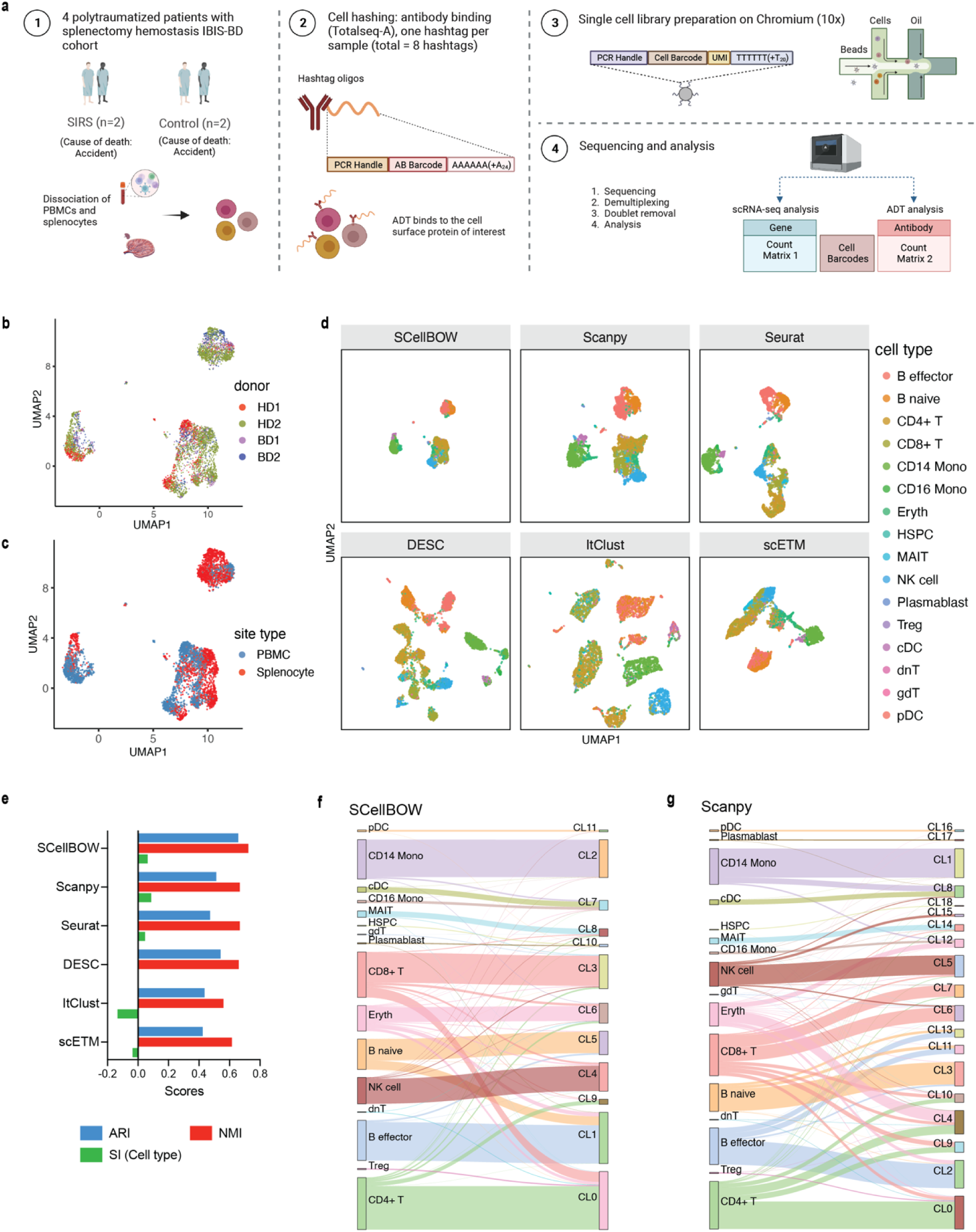
Evaluation of in-house splenocytes and matched PBMC dataset. **a,** An experiment schematic diagram highlighting the sites of the organs for tissue collection and sample processing. In this matched PBMC-splenocyte CITE-seq experiment, PBMCs and splenocytes were collected, followed by high-throughput sequencing and downstream analyses. **b-c,** UMAP plots for SCellBOW embedding colored by donors (b) and cell types (c). **d,** The UMAP plots for the embedding of SCellBOW compared to different benchmarking methods. The coordinates of all the plots are colored by cell type annotation results using Azimuth. **e,** Bar plot for ARI, NMI, cell type SI at resolution 1.0. **f-g,** Alluvial plots for Azimuth cell types mapped to SCellBOW clusters (f) and Scanpy clusters (g). The resolution of SCellBOW was set to 1.0. CL is used as an abbreviation for cluster.

### SCellBOW facilitates survival-risk attribution of tumor subpopulations

Every cancer features unique genotypic as well as phenotypic diversity, impeding the personalized management of the disease. The widespread adoption of genomics in cancer care has allowed correlating molecular portraits of tumors with patient survival in all major cancers. A few studies suggested that there is an association between the tumorigenicity of stromal cells in tumor microenvironments and patient survival^43^. This has sparked debate about the utility of single-cell gene expression in profiling single tumor cells. SCellBOW’s *phenotype algebra* module estimates the relative aggressiveness of different malignant and non-malignant cell subtypes using survival information as a surrogate. This presents an opportunity to gauge the aggressiveness of each cancer cell subtype in a tumor. For example, subtracting an aggressive phenotype from the *total tumor* (average of SCellBOW embeddings across all cells in a tumor) would better the odds of survival relative to dropping a subtype under negative selection pressure. By associating the embedding vectors, representing *total tumor* – *a specific cell cluster,* with tumor aggressiveness, *phenotype algebra* immediately opens a way to infer the level of aggressiveness of a particular cluster of cancer cells obtained through single-cell clustering.

As proof of concept, we first validated our approach on glioblastoma multiforme (GBM), which has been studied widely employing single-cell technologies. GBM has three well-characterized malignant subtypes: proneural (PRO), classical (CLA), and mesenchymal (MES)^44,45^. We obtained known markers of PRO, CLA, and MES to annotate 4,508 malignant GBM cells obtained from a single patient reported by Couturier and colleagues^46^ and used it as our target data (**Fig. 4a, Supplementary Methods 2.1**). As the source data, we used the GBM scRNA-seq data from Neftel *et al.*^47^. The survival data consisted of 613 bulk GBM samples with paired survival information from the TCGA consortium. We constructed the following query vectors for the survival prediction task: *total tumor*, i.e., average of embeddings of all malignant cells, *total tumor* – (MES+CLA), *total tumor* – MES/CLA/PRO (individually). We conjectured that for the most aggressive malignant cell subtype, *total tumor – subtype specific pseudo-bulk*, would yield the biggest drop in survival risk relative to the *total tumor*. Survival risk predictions associated with *total tumor* – MES/CLA/PRO thus obtained reaffirmed the clinically known aggressiveness order, i.e., CLA MES PRO, where CLA succeeds the rest of the subtypes in aggressiveness^48^ (**Fig. 4c, d**). More complex queries can be formulated, such as *total tumor* – (MES+CLA), which indicates that the tumor does not comprise the two most aggressive phenotypes, CLA and MES, and instead consists only of PRO cells. Hypothetically, this represents the most favorable scenario, as testified by the *phenotype algebra*^45,49^.

**Fig. 4.**
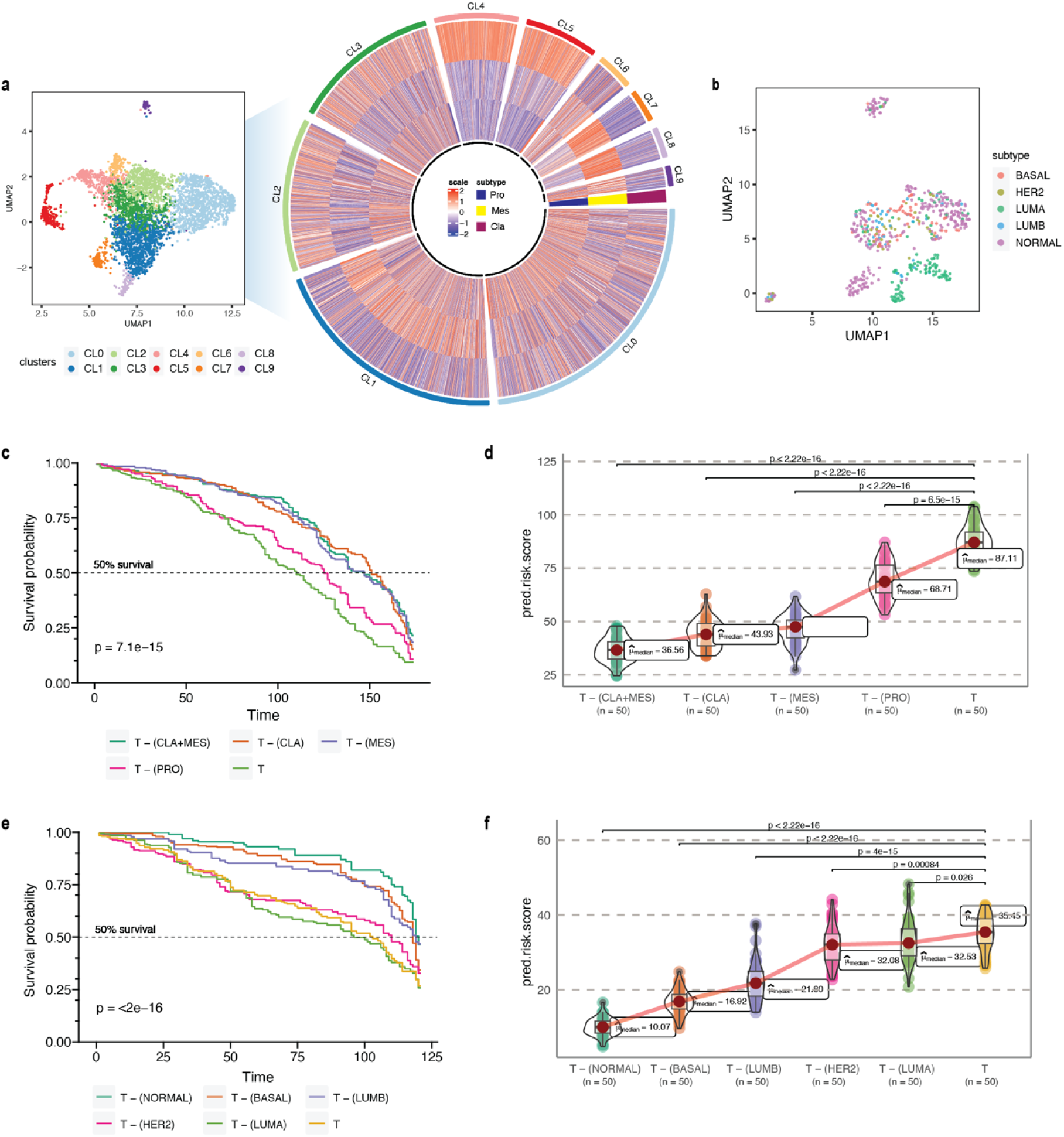
*Phenotype algebra* on GBM and BRCA known molecular subtypes. **a,** Heatmap for GSVA score for three molecular subtypes of GBM: CLA, MES, and PRO, grouped by SCellBOW clusters at resolution 1.0. **b,** UMAP plot for the embedding of BRCA target dataset colored by PAM50 molecular subtype. **c,** Survival plot for GBM molecular subtypes based on *phenotype algebra*. **d,** Violin plot for predicted risk scores for GBM molecular subtypes. **e,** Survival plot for BRCA molecular subtypes based on *phenotype algebra*. The *total tumor* is denoted by *T*. **f,** Violin plot for predicted risk scores for BRCA molecular subtypes.

We performed a similar benchmarking on well-established PAM50-based^50^ breast cancer (BRCA) subtypes: luminal A (LUMA), luminal B (LUMB), HER2-enriched (HER2), basal-like (BASAL), and normal-like (NORMAL) (**Fig. 4b**). We used 24,271 cancer cells from Wu *et al.*^51^ as the source data, 545 single-cell samples from Zhou *et al.*^52,53^ as the target data. We used 1,079 bulk BRCA samples with paired survival information from TCGA as the survival data. SCellBOW-predicted survival risks for the different subtypes were generally in agreement with the clinical grading of the PAM50 subtypes^54,55^. Exclusion of LUMA from *total tumor* yielded the highest risk score, indicating that LUMA has the best prognosis, followed by HER2 and LUMB, whereas BASAL and NORMAL were assigned worse prognosis^56–58^ (**Fig. 4e, f**). We observed an interesting misalignment from the general perception about the relative aggressiveness of the NORMAL subtype-removal of this subtype from *total tumor* indicated the highest improvement in prognosis. The NORMAL subtype is a poorly characterized and rather heterogeneous category. Recent evidence suggests that these tumors potentially represent an aggressive molecular subtype and are often associated with highly aggressive claudin-low tumors^54,59^. These benchmarking studies on the well-characterized cancer subtypes of GBM and BRCA affirm SCellBOW’s capability to preserve the desirable characteristics in the resulting phenotypes obtained as outputs of algebraic expressions involving other independent phenotypes and operators such as +/–. To further validate the efficiency of SCellBOW in learning single-cell latent representation, we have illustrated the effectiveness of fixed-length embeddings obtained from SCellBOW for *phenotype algebra* compared to other methods-scETM, scBERT, and scPhere (**Supplementary Fig. 6**, **Fig. 5f, and Supplementary Note 4**). Our results showed that SCellBOW learned latent representation of single-cells accurately captures the ‘semantics’ associated with cellular phenotypes and allows algebraic operations such as ‘+’ and ‘–’. Whereas scETM, scBERT, and scPhere fail to stratify cancer clones in terms of their aggressiveness and contribution to disease prognosis.

**Fig. 5.**
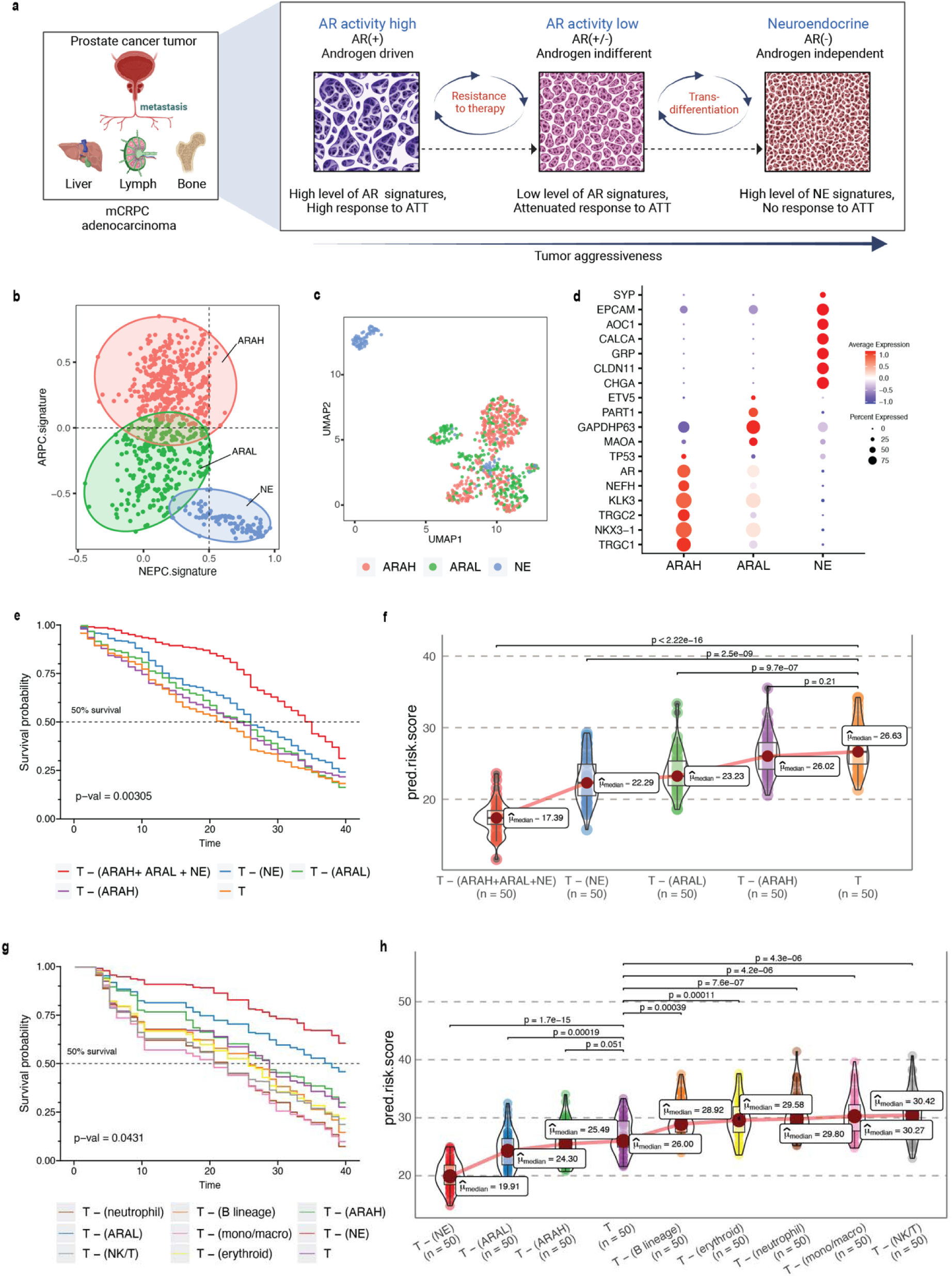
*Phenotype algebra* on mCRPC known molecular subtypes based on AR- and NE-activity. **a,** Schematic of the transdifferentiation states underlying lineage plasticity that occurs during mCRPC progression from an ARPC to NEPC. **b,** Scatter plot of GSVA scores of ARPC and NEPC gene sets, K-means clustering was used to allocate cells into the three high-level ARAH, ARAL, and NEPC categories. **c,** UMAP plot for projection of SCellBOW embedding colored by ARAH, ARAL, and NEPC. **d,** Heatmap showing the top differentially expressed genes (y-axis) between each high-level category (x-axis) and all other cells, tested with a Wilcoxon rank-sum test. **e,** Survival plot for mCRPC cancer phenotypes based on *phenotype algebra*. The t*otal tumor* is denoted by *T*. **f,** Violin plot for predicted risk scores for mCRPC phenotypes - ARAH, ARAL, and NEPC. **g,** Survival plot for mCRPC tumor microenvironment phenotypes based on *phenotype algebra*. The t*otal tumor* is denoted by *T*. **h,** Violin plot for predicted risk scores for mCRPC tumor microenvironment phenotypes - ARAH, ARAL, and NEPC.

### SCellBOW enables stratification of prostate cancer tumor microenvironment

In prostate cancer, the processes of transdifferentiation and dedifferentiation are vital in metastasis and treatment resistance^60^ **(Fig. 5a)**. Prostate cancer originates from secretory prostate epithelial cells, where AR, a transcription factor regulated by androgen, plays a key role in driving the differentiation^61,62^. Androgen-targeted therapies (ATTs) constitute the primary treatment options for metastatic prostate cancer, and they are most effective in well-differentiated prostate cancer cells with high AR activity^63^. After prolonged treatment with ATTs, the cancers eventually progress towards metastatic castration-resistant prostate cancer (mCRPC), which is highly recalcitrant to therapy^64^. In response to more potent ATTs, the prostate cancer cells adapt to escape reliance on AR with low AR activity^65^. The loss of differentiation pressure results in altered states of lineage plasticity in prostate tumors^66,67^. The most well-defined form of treatment-induced plasticity is neuroendocrine transformation. NEPC is highly aggressive, that often manifests with visceral metastases and currently lacks effective therapeutic options^66,68^. Recent studies have pointed toward the existence of additional prostate cancer phenotypes that emerge through lineage plasticity and metastasis of malignant cells^14^. This includes malignant phenotypes such as low AR signaling and DNPC, which lack AR activity and NE features. These additional phenotypes, resulting from the mechanisms of resistance to AR inhibition, can likewise be characterized by distinct gene expression patterns. Presumably, these phenotypes represent an intermediate or transitory state of the progression trajectory from high AR activity to neuroendocrine transdifferentiation^69^.

Here, we performed a pooled analysis of scRNA-seq target data consisting of 836 malignant cells derived from tumors collected from 11 tumors (mCRPC)^70^. We initially classified cells into three categories - AR activity high (AR+/NE–, ARAH), AR activity low (AR_low_/NE–, ARAL), and neuroendocrine (AR–/NE+, NE). The classification was based on the known molecular signatures associated with ARPC and NEPC genes^69^ (**Fig. 5b, Supplementary Methods 2.2**). We used the pre-trained model built on the Karthaus *et al.*^32^ dataset for transfer learning. We used 81 advanced metastatic prostate cancer patient samples with paired survival information from Abida *et al.*^23^ as the survival data. We subsequently assessed the relative aggressiveness of these high-level categories using *phenotype algebra* (**Fig. 5e, f**). We observed that subtracting the latent signature associated with the NE subtype from the tumor led to the largest drop in the predicted risk score, aligning with the anticipated order of survival^71^. In contrast, removing the ARAH subtype from the *total tumor* had a minimal impact on the predicted risk score. Furthermore, subtracting the ARAL subtype resulted in a predicted risk score between ARAH and NE, which supports the hypothesis that ARAL represents an intermediate state between ARAH and NE^69^. Upon further examination of the tumor prognosis by removing the subtypes in a combined state (ARAH+ARAL+NE), the *total tumor* had the highest positive improvement. The survival probability graph also followed the same order.

Till now, *phenotype algebra* has been used primarily for the stratification of cancer subpopulations. The tumor microenvironment includes malignant cells as well as various non-malignant cell types. This diverse cellular composition significantly influences the response to drug treatments. While malignant cells within a tumor typically show a great extent of heterogeneity, other cell types, such as immune cells, fibroblasts, and others, go through functional changes and variations in the amount of tumor infiltration. Therefore, it is crucial to distinguish and analyze the expression signals originating from the tumor cells in order to get a clearer picture of the gene expression changes in the cancer cells. This approach allows for a more accurate characterization of cancer subtypes based on the intrinsic properties of the malignant cells themselves. However, it is not always straightforward to unambiguously distinguish malignant from non-malignant cells in the complex environment of a tumor. To evaluate the capability of *phenotype algebra* in distinguishing between malignant and non-malignant cells, we applied it to single-cell transcriptome profiles of the mCRPC tumor microenvironment. This dataset, obtained from He et al., consists of 2,170 cells, including malignant cells as well as various immune and stromal cell types (**Supplementary Fig. 8h)**. We used the authors’ annotations for cell-type classification. We observed that the predicted risk score decreases when aggressive cancer subtypes are removed from the whole tumor population (**Fig. 5g, h**). Conversely, the risk score increases when immune cells are removed, suggesting that immune presence influences overall tumor aggressiveness. These results highlight the ability of *phenotype algebra* to capture and quantify risk signals from gene expression data, effectively distinguishing between malignant and non-malignant subpopulations in the tumor microenvironment.

### Unsupervised risk-stratification of metastatic prostate cancer clusters using SCellBOW

Grouping prostate cancer cells into three high-level categories is an oversimplified view of the actual heterogeneity of advanced prostate cancer biology. Herein, SCellBOW clusters could be utilized to discover novel subpopulations based on the single-cell expression profiles that result from therapy-induced lineage plasticity. Subsequently, *phenotype algebra* can assign a relative rank to these clusters under a negative selection pressure based on their aggressiveness. We utilized this concept to cluster the malignant cells from He *et al.* using SCellBOW, resulting in eight clusters, and then we predicted the relative risk for each cluster (**Fig. 6a, b**). This approach enables a novel and more refined understanding of lineage plasticity states and characteristics that determine aggressiveness during prostate cancer progression. This goes beyond the conventional categorization into ARAH, ARAL, and NE.

**Fig. 6.**
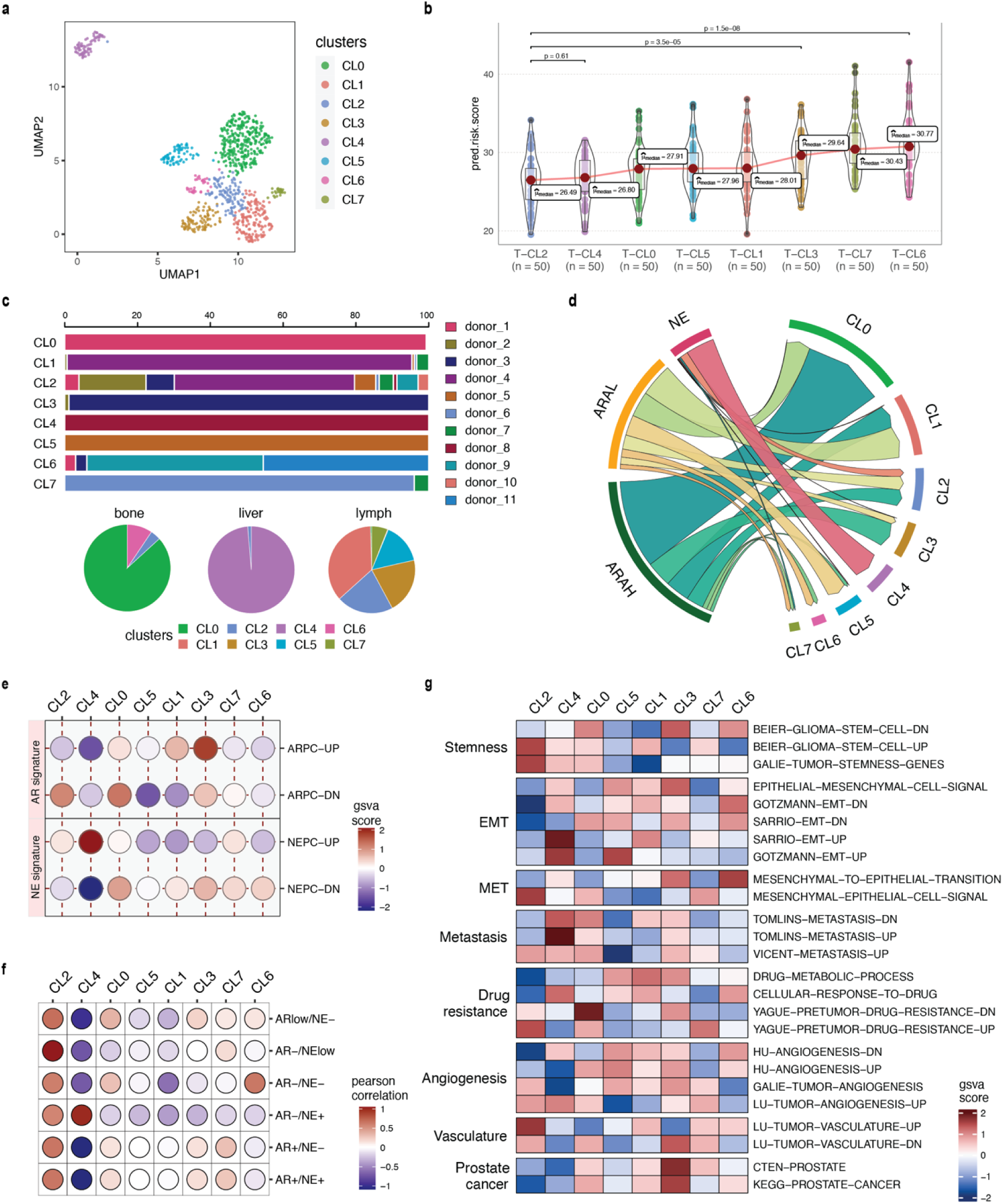
*Phenotype algebra* on He *et al.*^70^ mCRPC data based on SCellBOW clusters. **a,** UMAP plot for projection of embeddings with coloring based on the SCellBOW clusters at resolution 0.8. CL is used as an abbreviation for cluster. **b,** Violin plot of *phenotype algebra*-based cluster-wise risk scores for SCellBOW clusters based on *phenotype algebra*-based predictions. **c,** Patient and organ site distribution across the SCellBOW clusters. **d,** Illustration of the distribution of cells from the three high-level groups-ARAH, ARAL, and NEPC across the SCellBOW clusters. **e,** Bubble plot of row-scaled GSVA scores for custom curated gene sets containing activated and repressed AR- and NE-signatures. **f,** Correlation plot of six phenotypic categories based on DSP gene expression correlated with the SCellBOW clusters based on scRNA-seq gene expression. The six phenotypic categories are defined by Brady *et al.* based on the activity of AR and NE programs. **g,** Top gene sets correlated with SCellBOW clusters. Signatures were collected from the C2 ‘‘curated’’, C5 ‘‘Gene Ontology’’, and H ‘‘hallmark’’ gene sets from mSigDB^94^. Ranking by row scaled GSVA scores of one cluster against all others.

To further elucidate these altered cellular programs, we performed a gene set variation analysis (GSVA)^72^ based on the AR- and NE-activity (**Supplementary Methods 3, and Supplementary Data 1**). Our result showed that CL4 is characterized by the highest expression of NE-associated genes and the absence of AR-regulated genes, indicating the conventional NEPC subtype. Despite CL4 having the strongest NEPC signatures, eliminating CL2 from the tumor conferred an even higher aggressiveness level. Overall, we observed that, unlike other clusters, CL2 is composed of cells from the majority of drug-treated patients and multiple metastatic sites (**Fig. 6c**). This highlights that the clustering is not confounded by the individuals or the tissue origin, as often observed during integrative analysis of tumor scRNA-seq data. CL2 instead features a unique gene expression profile common to these cells. Moreover, CL2 has a mixed signature entailing ARAH, ARAL, and NE, indicating the emergence of a more transdifferentiated subtype as a consequence of therapy-induced lineage plasticity (**Fig. 6d**). Even though CL2 shows NE signature, it is distinguished by the gene expression signature induced by the inactivation of the androgen signaling pathway due to ATTs. As a consequence, cells manifesting this novel signature are grouped into a single cluster (**Fig. 6e**). Among all clusters, CL1 and CL3 resemble the traditional ARAH subtype. According to our *phenotype algebra* model, excluding CL6 and CL7 from the *total tumor* yielded the highest risk score. Brady *et al.*^13^ have broadly partitioned metastatic prostate cancer into six phenotypic categories using digital spatial profiling (DSP) transcript and protein abundance data in spatially defined metastasis regions. To categorize the SCellBOW clusters into these broad phenotypes, we performed Pearson’s correlation test between averaged DSP expression measurements of the six phenotypes and averaged scRNA-seq expression of the SCellBOW clusters (**Fig. 6f**). The results revealed that CL2 has the highest correlation with the phenotype defined by lack of expression of AR signature genes and low or heterogeneous expression of NE-associated genes (AR–/NE_low_). Similarly, as expected, CL4 showed the highest correlation with the NEPC phenotype defined by positive expression of NE-associated genes without AR activity (AR–/NE+). Meanwhile, CL6 exhibits a closer resemblance to a DNPC phenotype (AR–/NE–).

To gain a deeper understanding of these modified cellular processes in CL2 compared to other clusters, we conducted cluster-wise functional GSVA based on the hallmarks of cancer (**Fig 6g**). We observed that CL2 exhibited the least prostate cancer signature, indicating that this cluster has deviated from prototypical prostate cancer behavior. It has rather dedifferentiated into a more aggressive phenotype as a consequence of therapy-induced lineage plasticity^73^. CL2 exhibited the highest enrichment of genes related to cancer stemness compared to other clusters. CL2 showed pronounced repression of epithelial genes that are downregulated during the epithelial-to-mesenchymal transition (EMT). Furthermore, there is a lack of expression of genes that are upregulated during the reversion of mesenchymal to epithelial phenotype (MET). Thus, the cells in CL2 are undergoing a process of dedifferentiation from being epithelial cells and activating the mesenchymal gene networks. Existing studies have reported that adaptive resistance is positively correlated with the acquisition of mesenchymal traits in cancer^74,75^. CL2 was enriched with signatures associated with metastasis and drug resistance. Specifically, CL2 showed downregulation of the genes involved in drug metabolism and cellular response while upregulation of the genes associated with drug resistance. In cancer cells, the acquisition of stemness-like as well as cell dedifferentiation (mesenchymal) traits can facilitate the formation of metastases and lead to the development of drug resistance^76^. Further analysis of the metastatic potential of CL2 indicated that the cluster is enriched with genes associated with vasculature and angiogenesis. Tumors induce angiogenesis in the veins and capillaries of the host tissue to become vascularized, which is crucial for their growth and metastasis^77^. Thus, based on our findings, we anticipate that CL2 corresponds to a highly aggressive and dedifferentiated subpopulation of mCRPC within the lineage plasticity continuum, correlating to poor patient survival and positive metastatic status. Our cluster-wise *phenotype algebra* results imply that the traits of the androgen and neuroendocrine signaling axes are not the exclusive defining features of the predicted risk ranking and that other yet under-explored biological programs play additional important roles.

## OVERALL DISCUSSION

In this work SCellBOW, a scRNA-seq analysis framework inspired by NLP-based transfer learning approach can be utilized to decipher molecular heterogeneity from scRNA-seq profiles and infer survival risks associated with the individual cell subpopulations in tumor microenvironment. In this novel approach, cells are treated as a bag-of-molecules, similar to representing a document as a “bag” of words (BOW), representing molecules in proportion to their abundance within each cell. SCellBOW uses a document embedding architecture to preserve the cellular similarities observed in the gene expression space. SCellBOW-learned neuronal weights are transferable. The model can use the knowledge acquired from a source dataset to warm start the learning process on a target dataset. Although Doc2vec is not commonly used for transfer learning, our experiments, which involve repurposing a pre-trained, modified Doc2vec model for scRNA-seq analysis, establish promising use cases for transfer learning (**Supplementary Fig. 7 and Supplementary Note 5**). SCellBOW accomplishes two major tasks: a. single-cell representation learning under a transfer learning framework facilitating high-quality cell clustering and visualization of scRNA-seq profiles. b. *phenotype algebra*, enabling the attribution of survival risk to tumor cell clusters obtained from tumor scRNA-seq data. For both tasks, we compared SCellBOW with several state-of-the-art single-cell analysis methods, including ones based on complex language modeling architectures (e.g., scETM uses a topic model, scBERT uses a transformer-inspired model). SCellBOW exhibited consistency in characterizing cellular heterogeneity through clustering and survival risk (aggressiveness) stratification of tumor cell subtypes.

With the SCellBOW, we observe considerable improvement in the quality of clustering and *phenotype algebra* results obtained from scRNA-seq target datasets. The existing single-cell transfer learning methods, such as scETM, ItClust, and scBERT, are pre-trained on large amounts of pre-annotated single-cell data to achieve good performance. However, using source data annotations as a reference limits the identification of new cell types in the target data. Moreover, the degree of imbalance in the cell type distribution substantially influences the performance of these methods^78^. It is important to realize that exploratory studies involving single-cell expression profiling require unsupervised data analysis. SCellBOW addresses these challenges by allowing self-supervised pre-training on gene expression datasets, learning the general syntax and semantic patterns from the unlabeled scRNA-seq dataset. Additionally, it is less susceptible to overfitting^79^, especially with small training datasets, owing to the shallow neural network used in SCellBOW, which enables faster convergence compared to other studied transfer learning methods (**Supplementary Table 5**). Notably, SCellBOW, unlike ItClust and scETM, does not limit model training to genes intersecting between the source and the target datasets and is independent of the source data cell type annotation. Moreover, while methods such as scETM, ItClust, and scBERT utilize numerical gene expression values directly as an input to their network architectures, SCellBOW, for the first time, encodes gene expression profiles into documents in their native format.

We compared the clustering capability of SCellBOW to popular single-cell analysis techniques as well as some recently proposed transfer learning approaches. An array of scRNA-seq datasets was used for the same, representing different sizes, cell types, batches, and diseases. As evident from the comparative analyses, SCellBOW exhibits robustness and consistency across all data and metrics. We also reported tangible benefits of transfer learning from apt source data. For example, transfer learning with a large number of tumor-adjacent normal cells from the prostate as the source and healthy prostate cells as the target adequately portrayed the cell type diversity therein. SCellBOW reliably clustered pancreatic islet-specific cells that were processed using varied single-cell technologies. The prevalence of single-cell expression profiling and the heterogeneity of PBMCs make it one of the best-studied tissue systems in humans^80^. We isolated matched PBMCs and splenocytes from two healthy donors and two brain-dead donors (the cause of death is subarachnoid hemorrhage on aneurysm rupture). The utilization of this data presents a more intricate problem, where the cells originate from diverse biological and technical replicates, conditions, individuals, and organs. SCellBOW-based analysis of PBMCs with near-ground-truth cell type annotation confirmed its ability to adequately decipher underlying cellular heterogeneity. Our results allude to a visible improvement in scRNA-seq analysis outcomes, even with small sample sizes and multiple covariates, when contrasted with other best-practice single-cell clustering methods.

Beyond robust identification of cell type clusters, SCellBOW uses algebraic operations to analyze the cellular phenotypes that could potentially identify their contribution to patient survival outcomes. We have leveraged the power of word algebra through document embeddings to perform risk stratification of cancer subtypes based on their aggressiveness. This involves comparing the likelihood of a subtype being eliminated from the entire tumor under negative selection pressure. In addition to overall survival probability, SCellBOW assigns a risk score to discern the differences between equally aggressive subpopulations that may be hard to decipher from their survivability profiles. The *phenotype algebra* module exhibits resilience in describing the risk associated with malignant cell subpopulations arising from various types of cancer, which otherwise cannot be accomplished using gene expression data (**Supplementary Note 7, Supplementary Fig. 8i-k**). We demonstrated a potential use case in estimating the survival risk of known molecular subtypes of three cancer types: GBM, BRCA, and mCRPC. Several examples were shown where simple algebraic operators such as ‘+’ and ‘–’ could derive clinically intuitive outcomes. For example, subtracting the CLA phenotype from the whole tumor resulted in an improvement in the survival risk in a GBM patient compared to MES and PRO subtypes. Upon further probing of the tumor prognosis by removal of the subtypes in combinations from the tumor, specifically, CLA and MES subtypes, which are known to be the most aggressive, we observed the highest improvement. To summarize, SCellBOW allows simulations involving multiple phenotypes. Our subsequent investigation focused on a BRCA dataset that harbors a more complex subtype structure. We observed a deviance in our results from the general perception about the relative aggressiveness of the NORMAL subtype in breast cancer. Notably, removing the NORMAL subtype from the *total tumor* was associated with the highest improvement in prognosis. This contradicts the common assumption that a NORMAL subtype is an artifact resulting from a high proportion of normal cells in the tumor specimen^81^. Despite indications that these tumors often do not respond to neoadjuvant chemotherapy^58^, the clinical significance of the NORMAL subtype remains uncertain due to a limited number of studies. The NORMAL breast cancer cells are poorly characterized and heterogeneous in nature. As per the recent classification of the breast cancer subtypes, the NORMAL subtype has been identified to be a potentially aggressive molecular subtype, referred to as claudin-low tumors^59^. This evidence suggests that the NORMAL subtype of breast cancer is potentially an aggressive molecular subtype, which is consistent with the prediction made by SCellBOW.

In advanced metastatic prostate cancer, molecular subtypes are still poorly defined, and characterizing novel or more fine-grained subtypes with clinical implications on patient prognosis is an active field of research. As a proof-of-concept, we executed SCellBOW on the three mCRPC subtypes, namely ARAH, ARAH, and NE. We observed that eliminating the subtypes individually and in combinations (ARAH+ARAL+NE) exhibited a clinically intuitive change in prognosis using SCellBOW. We further expanded the application of *phenotype algebra* beyond its previous application solely to cancer cells by including non-malignant cells from the tumor microenvironment. We observed that eliminating cancer subtypes from the tumor decreased its aggressiveness, whereas removing the immune component increased aggressiveness, highlighting the critical role of the tumor microenvironment in modulating cancer progression. To date, mCPRC classification has largely been confined to the gradients of AR and NE activities. There is considerable scope for embracing more fine-grained subtypes to better explain clonal selection and epithelial plasticity in drug resistance in wider use cases. Convinced by the overall performance of SCellBOW, we applied *phenotype algebra* on the SCellBOW clusters obtained from the He *et al.* mCRPC dataset^70^. We observed a misalignment from the general perception that the neuroendocrine subtype features the worst prognosis. Our results pointed towards the existence of a more aggressive and dedifferentiated subpopulation in the lineage plasticity continuum. This subpopulation of de-differentiated cells in mCRPC, AR−/NE_low_, exhibit greater aggressiveness and have distinct gene expression patterns that do not match those of any previously identified molecular subtypes. This novel subpopulation shares gene expression signatures with both androgen-repressed and neuroendocrine-activated genes. GSVA analysis of this hitherto unknown phenotype offered insights into its putative functional characteristics, which can be broadly defined by stemness, EMT, MET, and drug resistance. To summarize, SCellBOW clustering combined with *phenotype algebra* can provide novel insights into factettes of cancer biology. This can be used to delineate features of lineage plasticity and aid in better classification of molecular phenotypes with clinical significance. Once the subpopulations are identified, delving into the intricate mechanisms governing their resistant behavior can provide invaluable insights for designing new drugs. Further, such a tool can empower medical oncologists and oncology researchers to develop more personalized and systemic treatment regimes for cancer patients.

## CONCLUSION

SCellBOW is a scRNA-seq analysis tool that can be utilized to decipher molecular heterogeneity from scRNA-seq profiles and infer survival risks associated with the individual subpopulations in tumors. Our main contributions are as follows: a. We proposed SCellBOW which learns a distributed representation (fixed-length embedding per cell) of single cells. Despite being a computationally inexpensive and simple architecture, SCellBOW outperforms transformer-based approaches such as scBERT on both single-cell clustering and cancer cell risk stratification tasks. b. We also introduced the task of inferring cellular aggressiveness as a computationally tractable problem. The *phenotype algebra* module of SCellBOW performs joint representation learning of externally sourced tumor bulk RNA sequencing data (e.g., TCGA) and tumor scRNA-seq data of interest. Finally, a survival prediction model trained on bulk expression profiles is used for survival risk stratification for each of the malignant and non-malignant cell clusters derived from the tumor scRNA-seq data. We compared SCellBOW with existing best practice methods for its ability to precisely represent phenotypically divergent cell types across multiple scRNA-seq datasets, including an in-house generated human splenocyte and matched PBMC dataset. SCellBOW has proven effective in characterizing poorly defined metastatic prostate cancer. We identified a subpopulation of dedifferentiated cells in mCRPC, AR−/NElow, that exhibit greater aggressiveness and have distinct gene expression patterns that do not match those of any previously identified molecular subtypes. We could trace this back in a large-scale spatial omics atlas of 141 well-characterized metastatic prostate cancer samples at the spot resolution. In the case of breast cancer, our results indicated that the normal-like subtype, which was previously considered an artifact of high normal cell content in the tumor sample, may be among the most aggressive ones, which is concordant with recent reports. In conclusion, the robust clustering and risk-stratification capabilities of SCellBOW hold promise for tailored therapeutic interventions by identifying clinically relevant subpopulations and their impact on prognosis.

## MATERIAL AND METHODS

### Overview of SCellBOW components and functionalities

SCellBOW has two main applications: **a. *Cell embedding and clustering.*** In this work, we demonstratsingle-cell representation learning using a source and target scRNA-seq data. The source data is typically any large scRNA-seq data capable of priming a neural architecture for improved representation learning on the target data of interest. Note that in this case, cells in the source or the target data need not be annotated. This transfer learning capability is meant to obtain improved fixed-length embeddings for cells in the target data by leveraging the transcriptomic patterns from existing datasets. **b. *Survival risk attribution.*** SCellBOW’s *phenotype algebra module* predicts the aggressiveness associated with each malignant and non-malignant cell cluster. SCellBOW computes fixed-length embeddings jointly for the scRNA-seq target dataset (from the tumor under study) and tumor bulk RNA-seq profiles with patient-survival data (e.g., TCGA). Notably, embeddings of transcriptomes from scRNA-seq target data and bulk RNA-seq survival data have been determined concurrently by recalibrating the pre-trained model. A survival prediction model trained on the bulk RNA-seq embeddings and patient-survival data is then tested on the embeddings of the target data to make predictions about the degree of aggressiveness exhibited by the tumor variants. The steps involved in the above two tasks are depicted in Figure 1 (**Fig. 1**). The granular details involved in each of these steps can be found in the below subsections.

### Data preparation for clustering

SCellBOW follows the same preprocessing procedure for the source and target data. SCellBOW first filters the cells with less than 200 genes expressed and genes that are expressed in less than 20 cells. The thresholds may vary depending on the dimension of the dataset (**Supplementary Table 2**). After eliminating low-quality cells and genes, SCellBOW log-normalizes the gene expression data matrix. In the first step, CPM normalization is performed, where the expression of each gene in each cell is divided by the total gene expression of the cell, multiplied by 10,000. In the second step, the normalized expression matrix is natural-log transformed after adding one as a pseudo-count. Highly variable genes are selected using the highly_variable_genes() function from the Scanpy package. SCellBOW performs a z-score scaling on the log-normalized expression matrix with the selected highly variable genes. SCellBOW can handle data in different formats, including UMI count, FPKM, and TPM. The UMI count data follow the same preprocessing procedure as above. SCellBOW skips the normalization step for TPM and FPKM since their lengths have been normalized.

### Creating a corpus from source and target data

The gene expression data matrix is analogous to the term-frequency matrix, which represents the frequency of different words in a set of documents. In the context of genomic data, a similar concept can be used to represent the expression level of a gene (words) in a set of cells (documents). Let *E* ∈ *R^G^*^×*C*^ be the gene expression matrix obtained from a scRNA-seq experiment, where each value *E_g,c_* of the matrix indicates the expression value of a gene *g* ∈ *G* in a cell *c* ∈ *C* obtained after Scanpy preprocessing. SCellBOW generates embeddings by taking two input datasets: a source and a target data matrix. The source dataset contains the initial weights of the neural network model, while the target data contains the cells that require clustering or malignancy potential ranking. Before generating the embeddings, SCellBOW performs feature scaling on the data matrix, rescaling each feature to a range of [0, 10] as follows.

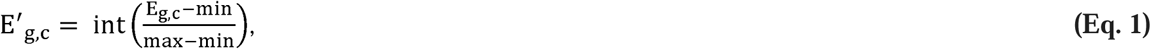

where scaling is applied to each entry with *max* = 10 and *min* = 0. This establishes the equivalence between term-frequency and gene expression matrix. Here, we consider that a specific gene is a word, and the expression of the gene in a cell corresponds to the number of copies of the word in a document. To scale the data matrix, we have used the MinMaxScaler() function from the scikit-learn package^82^. To create the corpus, SCellBOW duplicates the name of the expressed gene in a cell as many times as the gene is expressed. SCellBOW shuffles the genes in each cell to ensure a uniform distribution of genes across the cell, removing any positional bias within the dataset^19^. We verified that this randomization does not change the outcomes dramatically (**Supplementary Fig. 9**). The resulting gene names are treated as the words in the document. To build the vocabulary from a sequence of cells, a tag number is associated with each cell using the TaggedDocument() function in the Gensim package^83^. For each gene in the documents, a token is assigned using a module called tokenize with a word_tokenize() function in NLTK package^84^ that splits gene names into tokens.

### The SCellBOW network

SCellBOW produces a low-dimensional fixed-length embedding of the single-cell transcriptome using transfer learning. The data matrix is transformed from E′ ∈ *Z^G^*^×*C*^ feature space to E″ ∈ *R^d^*^×*C*^ latent feature space of d dimensions, with *d* ≪ *G*. To generate the embeddings, SCellBOW trains a Doc2vec distributed memory model of paragraph vectors (PV-DM) model. The PV-DM model is similar to bag-of-words models in Word2vec^85^. The training corpus in Doc2vec contains a set of documents, each containing a sequence of words *W* = {*w*_1_, *w*_2_, …, *w*_T_} that forms a vocabulary *V*. The words within each document are treated as shared among all documents. The training objective of the model is to maximize the probability of predicting the target word *w_t_*, given the context words that occur within a fixed-size window of size *n* around *w_t_* in the whole corpus as follows.

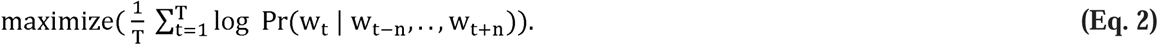

The probability can be modeled using the hierarchical SoftMax function as follows.

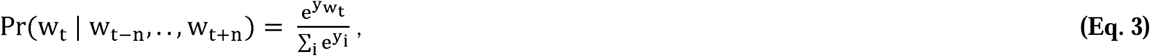

where each of *y_i_* computes the log probability of the word *w_t_* normalized by the sum of the log probabilities of all words in V. This is achieved by adjusting the weights of the hidden layer of a neural network. To build the Doc2vec model, we used the doc2vec() function in the Gensim library. The initial learning rate was set to 0.025, and the window size to 5. We chose the PV-DM training algorithm and set the embedding vector size to 300 as the default parameter. The choice of embedding vector size may be adjusted according to the size and dimensionality of the dataset.

### Fine-tuning using transfer learning

At first, SCellBOW is pre-trained with a source data matrix 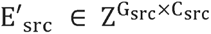 genes and G_src_ cells. The source data matrix sets the initial weights of the neural network model. During transfer learning, SCellBOW fine-tunes the weights learned by the pre-trained model from *E*′*_src_* using a target dataset 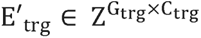 with G_trg_ genes and C_trg_ cells. This facilitates faster convergence of the neural network compared to starting from randomly initialized weights. The output layer of the network is a fixed-length low-dimension embedding (a vector representation) for each cell c ∈ C_trg_. To infer the latent structure of *E_trg_* single-cell corpus, we used the infer_vector() function in the Gensim package to produce the dimension-reduced vectors for each cell in C_trg_. This step ensures the network can map the target data into a low dimensional embedding space *R^d^*; i.e., E^’^→E″, where E″ ∈ R^d×C^. The resulting embeddings can be used for various downstream analyses of the single-cell data, such as clustering, visualization of cell types, and *phenotype algebra*.

### Visualization of SCellBOW clusters

SCellBOW maps the cells to low-dimensional vectors in such a way that two cells with similar gene expression patterns will have the least cosine distance between their inferred vectors.

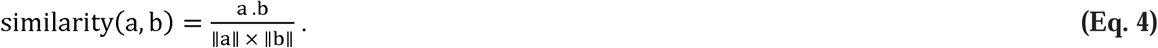

After generating low-dimensional embeddings for the cells, SCellBOW identifies the groups of cells with similar gene expression patterns. To determine the clusters, SCellBOW uses the Leiden algorithm^86^ on the embedding matrix of the target dataset. We used the leiden() function in the Scanpy package with a default resolution of 1.0. The resolution might vary in the reported results depending on the dimension of the target dataset. To visualize the clusters within a two-dimensional space, SCellBOW uses the umap() function from the Scanpy library.

### Data preparation for *phenotype algebra*

Three independent datasets are required to perform *phenotype algebra*. Two of the three datasets are scRNA-seq datasets used for transfer learning (source and target data). SCellBOW preprocesses the source data matrix using the standard preprocessing steps. SCellBOW uses an additional bulk RNA-seq gene expression matrix (referred to as survival data). The samples are paired with survival information (e.g., vital status, days to follow-up, days to death). In the survival data, samples without follow-up time or survival status and samples with clinical information but no corresponding RNA-seq data were excluded. SCellBOW accounts for unequal cell distribution across different classes in the target data matrix. SCellBOW up samples the imbalanced target dataset by generating synthetic samples from the minority class. The value of the synthetic minority class sample is determined by interpolating between its neighboring cells from the same class. We used SMOTE^87^ from the imblearn python library^88^. This confirms that the cell type proportions do not confound the output of *phenotype algebra*.

### Generating pseudo-bulk vectors for algebraic operations

SCellBOW constructs two types of pseudo-bulk vectors from the target dataset. First, it creates a pseudo-bulk reference vector by averaging the gene expression of all cells in the tumor (*total tumor*). This serves as a baseline representation of the entire tumor and acts as a reference for ranking different groups based on their relative expression profiles. Next, for each group in the dataset, SCellBOW generates a pseudo-bulk phenotype vector by averaging the gene expression of the cells within that specific group. This allows for a direct comparison between different cellular phenotypes. The phenotype vectors are constructed either based on the user-defined cell populations or SCellBOW clusters. Additionally, SCellBOW can perform algebraic operations, such as addition or subtraction, on multiple phenotype vectors to assess the combined risk of two or more cellular phenotypes. For example, in the equation (P_a_ + P_b_), P_a_ and P_b_ represents the phenotype vectors of two distinct cellular phenotypes. Following this, SCellBOW concatenates the bulk RNA-seq data matrix, and the reference and phenotype vectors based on common genes. We used the concatenate() function in the AnnData python package^89^. The resulting combined dataset is then passed to the pre-trained model, which maps it to a lower-dimensional embedding space.

### Survival risk attribution

SCellBOW infers the survival probability and predicted risk score for the user-defined tumor subtypes and SCellBOW clusters. The survival risk of the different groups is predicted by fitting a random survival forest (RSF)^90^ machine learning model with SCellBOW embeddings. At first, the RSF is trained with the survival information combined with bulk RNA-seq embeddings obtained from transfer learning. During the prediction step, pseudo-bulk embeddings generated from the target dataset, serving as the test data, undergo a subtraction operation in which the cosine distance between each query embedding and the reference embedding is calculated. For example, the equations:

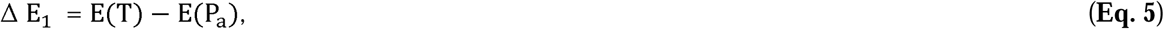

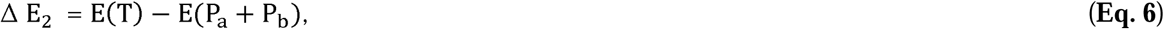

where E(T) represents the embedding of the average expression for the total tumor, while E(P_a_) and E(P_a_ + P_b_) represent the embeddings of the phenotype vectors, individually and in combination, respectively. Here, Δ *E*_1_ and Δ *E*_2_ represent the differences between the embeddings, simulating the removal of subtypes from the whole tumor under negative selection pressure. The resulting difference is then used as input to the RSF model to infer the survival probability S(t). The survival probability computes the probability of occurrence of an event beyond a given time point *t* as follows.

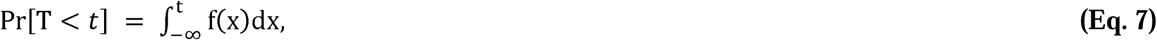

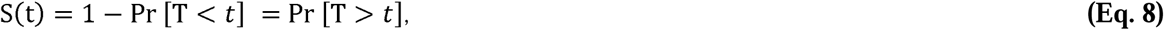

where *T* denotes the waiting time until the event occurs and *f*(*x*) is the probability density function for the occurrence of an event. SCellBOW computes the survival probability using the predict_survival_function() from the scikit-survival RandomSurvivalForest python package^91^ with n_estimators = 1000.

In addition to survival probability, SCellBOW can estimate the relative aggressiveness of different phenotypes by assigning a risk score for distinct groups. To infer the predicted risk score, SCellBOW first trains 50 bootstrapped RSF models using 80% of the training set for each iteration. The training data is sub-sampled using different seeds for every iteration. We used the predict() function from the scikit-survival package to compute the risk score of each of the input vectors. SCellBOW derives the median of the predicted risk score for each group from the 50 bootstrapped models. The *phenotype algebra* model assigns groups with shorter survival times a lower rank by considering all possible pairs of groups in the data. The groups with a lower predicted risk score after removal from the reference pseudo-bulk are considered more aggressive, as they are associated with shorter survival times.

### Description of datasets for model evaluation

To evaluate the performance of SCellBOW, we used fifteen publicly available scRNA-seq datasets and an in-house scRNA-seq dataset spanning different cell types, sizes, and diseases (**Supplementary Table 2)**. Four use cases were constructed to benchmark the clustering efficiency, each involving a pair of single-cell expression datasets. In the first use case, we used ∼120,300 non-cancerous human prostate cells from Karthaus *et al.*^32^ and ∼28,600 healthy prostate cells from Henry *et al.*^33^. The second use case consisted of two PBMC datasets from Zheng *et al.*^34^ containing approximately 68,000 cells and 2,700 cells. For the third use case, we combined three independent batches of pancreatic islet cells from Baron *et al.*^35^, Muraro *et al.*^36^, and Wang *et al.*^37^, with a total of 11,181 cells as the source data and 2,068 cells from Segerstolpe *et al.*^38^ as the target data. In the fourth use case, we used the Zheng *et al.* data containing approximately 68,000 cells as the source data and the in-house dataset isolated from matched spleen and PBMC samples from multiple patients as the target data.

To validate the accuracy of *phenotype algebra*, we constructed three use cases from publicly available tumor datasets comprising cells from GBM, BRCA, and mCRPC patient tumors. Each instance involved a pair of single-cell expression datasets and a bulk RNA-seq dataset paired with clinical information. For ease of reference, we stick to the following nomenclature – a) the source data comprises the scRNA-seq dataset used for model pre-training, b) the target data comprises cells under investigation, and c) the survival data comprises tumor bulk RNA-seq samples. In the case of GBM, we used two scRNA-seq datasets comprising approximately 12,074 cells and 4,508 cells obtained from Neftel *et al.*^47^ and Couturier *et al.*^46^, respectively. In the case of BRCA, we retrieved triple-negative breast cancer scRNA-seq datasets from Wu *et al.*^51^ and Zhou *et al.*^53^ with approximately 24,271 cells and 545 malignant cells, respectively. In both the use cases, the bulk RNA-seq was obtained from TCGA (https://portal.gdc.cancer.gov). In the third use case, we acquired author-annotated malignant cells from He *et al.*^70^. A total of 836 malignant cells and 1334 non-malignant cells were derived from 11 patients and three metastasis sites: bone, lymph node, and liver. We retrieved 81 mCRPC bulk RNA-seq samples with paired survival from Abida *et al.*^23^. We obtained the gene expression of 141 regions of interest determined by digital spatial gene expression profiling of mCRPC from Brady *et al.*^13^. We used this data to correlate the He *et al.* scRNA-seq gene expression with the DSP gene expression of the six phenotypic categories based on the AR- and NE-activity: AR+/NE−, AR_low_/NE−, AR−/NE−, AR−/NE_low_, AR+/NE+, and AR−/NE+. Details of the datasets are described in **Supplementary Methods (Supplementary Methods 1)**.

### Isolation of matched PBMC and splenocyte samples

In this study, we isolated PBMC and splenocyte from four patients (two healthy donors, HD1 18 years old male and HD2 61 years old female; two brain-dead donors, BD1 57 years old female and BD2 58 years old female). PBMCs were isolated from blood after Ficoll gradient selection. Spleen tissue was mechanically dissociated and then digested with Collagenase and DNAseq. Splenocytes were finally isolated after Ficoll gradient selection. Splenocytes and PBMCs were stored in DMSO with 10% FBS in liquid nitrogen at the Biological Resource Centre for Biobanking (CHU Nantes, Hotel Dieu, Centre de Ressources Biologiques). This biocollection was authorized in May 2013 by the French Agence de la Biomedecine (PFS13-009).

### Library preparation

For the in-house dataset, we performed CITE-seq using Hashtag Oligos (HTO) to pool samples into a single 10X Genomics channel for scRNA-seq^92^. The HTO binding was performed following the specified protocol (Total-seq B). After thawing and cell washing, 1M cells were centrifuged and resuspended in 100uL PSE buffer (PBS/FBS/EDTA). Cells were incubated with 10ul human FcR blocking reagent for 10 min at 4°C. 1 uL of a Hashtag oligonucleotide (HTO) antibody (Biolegend) was then added to each sample and incubated at 4°C for 30 min. Cells were then washed in 1mL PSE, centrifuged at 500g for 5 min, and resuspended in 200uL PSE. Cells were stained with 2uL DAPI, 60uM filtered, and viable cells were sorted (ARIA). Cells were checked for counting and viability, then pooled and counted again. Cell viability was set to < 95%. Cells of HD1 spleen were lost during the washing step and thus not used during further downstream processing. Cells were then loaded on one channel of Chromium Next Controller with a 3’ single-cell Next v3 kit. We followed protocol CG000185, Rev C, until the library generation stage. For the HTO library, we followed the protocol of the 10X genomics 3’ feature barcode kit (PN-1000079) to generate HTO libraries.

### Next-generation sequencing and post-sequencing quality control

We sequenced 310pM of pooled libraries on a NOVAseq6000 instrument with an S1(v1) flow-cell. The program was run as follows: Read1 29 cycles / 8 cycles (i7) /0 (i5)/Read2 93 cycles (Standard module, paired-end, two lanes). The FASTQ files were demultiplexed with CellRanger v3.0.1 (10X Genomics) and aligned on the GRCh38 human reference genome. We recovered a total of 6,296 cells with CellRanger, and their gene expression matrices were loaded on R. We performed the downstream analysis of the in-house matched PBMC-splenocyte dataset in R using Seurat 4.0. For RNA and HTO quantification, we selected cell barcodes detected by both RNA and HTO. We demultiplexed the cells based on their HTO enrichment using the HTODemux() function in the Seurat R package with default parameters^27,93^. We subsequently eliminated doublet HTOs (maximum HTO count □>□1) and negative HTOs. Singlets were used for further analysis, leaving 4,819 cells and 33,538 genes. We annotated each of the cell barcodes using HTO classification as PBMC and Spleen based on the origin of cells and HD1, HD2, BD1, and BD2 based on the patients (**Supplementary Fig. 4**). We performed automatic cell annotation using the Seurat-based Azimuth using human PBMC as the reference atlas^42^ (**Supplementary Note. 2)**. We defined six major cell populations: B cells, CD4 T cells, CD8 T cells, natural killer (NK) cells, monocytes, and dendritic cells. We then manually verified the annotation based on the RNA expression of known marker genes (**Supplementary Fig. 4d and Supplementary Table 4**).

## AVAILABILITY OF DATA AND MATERIALS

The in-house matched PBMC-splenocyte scRNA-seq expression data generated in this study is available in the GEO with accession number GSE221007. Details of the public datasets analyzed in this paper are described in **Supplementary Table 2**.

All source codes are available at GitHub https://github.com/cellsemantics/SCellBOW.

## ACKNOWLEDGMENTS

DS acknowledges the support of the ihub-Anubhuti-iiitd Foundation set up under the NM-ICPS scheme of the DST. JP, AR, PS, and SF thank the biological resource centre for biobanking (CHU Nantes, Hôtel Dieu, Centre de Ressources Biologiques (CRB), Nantes, F-44093, France (BRIF: BB-0033-00040)) and the Genomics Core Facility GenoA, member of Biogenouest and France Genomique, and to the Bioinformatics Core Facility BiRD, member of Biogenouest and Institut Français de Bioinformatique (IFB) (ANR-11-INBS-0013) for the use of their resources and their technical support. Parts of the schematics were created with BioRender.com.

## AUTHOR CONTRIBUTIONS

DS conceived the study with CCN. JP, Antoine Roquilly, PS, and CF contributed to the experimental design of the matched PBMC-splenocyte CITE-seq study. NB developed the SCellBOW python package and conducted data analyses with assistance from Anja Rockstroh and SSD. SSD and SKT assisted NB in model training. DS supervised the algorithm development. CCN supervised cancer-related analyses and interpretation. Anja Rockstroh supervised and performed specific data analysis and data interpretation tasks. Anja Rockstroh, HK, JP, GA, ML, and BGH assisted in the biological interpretation of results. DS, CCN, and ML supervised the entire study. All authors substantially contributed to manuscript writing and reviewing.

