## Supplementary Information for "SCellBOW: AI-Driven Tumor Risk Stratification from Single-Cell Transcriptomics Using Phenotype Algebra"

**Affiliations:**

**Supplementary Table 1:** Overview of tools and benchmarking methods used in this paper

| **Tool** | **Version** | **URL** | **Resolution dependent** | **Visualization** | **Clustering algorithm** | **Reference** |
| --- | --- | --- | --- | --- | --- | --- |
| Scanpy | 1.9.1 | <https://github.com/scverse/scanpy> | Yes | UMAP | Louvain | Wolf *et al.**^1^* |
| ItClust | 1.2.0 | <https://github.com/jianhuupenn/ItClust> | No | UMAP | ItClust | Hu *et al.**^2^* |
| Seurat | 4.1.1 | <https://github.com/satijalab/seurat> | Yes | UMAP | Louvian | Butler *et al.**^3^* |
| DESC | 2.1.1 | <https://github.com/eleozzr/desc> | Yes | UMAP | Louvain | Li *et al.**^4^* |
| scBERT | 1.0.0 | https://github.com/TencentAILabHealthcare/scBERT | Yes | UMAP | Leiden | Yang *et al.**^5^* |
| scPhere | 1.0.0 | https://github.com/klarman-cell-observatory/scPhere | Yes | UMAP | Louvain | Ding and Regev *et al.^6^* |
| scETM | 0.4.9 | https://github.com/hui2000ji/scETM | Yes | UMAP | Leiden | Zhao *et al.**^7^* |

**Supplementary Table 2**: Summary of datasets analyzed in this paper

| **Model** | **Dataset** | **Tissue** | **Technology** | **Data Type** | **Cell/sample**  **Detected** | **Used in**  **SCellBOW** | **Data used**  **as** | **Cell filter** | **Gene filter** | **HVG** |
| --- | --- | --- | --- | --- | --- | --- | --- | --- | --- | --- |
| Normal Prostate | Karthaus *et al.**^8^* | Human primary  prostate cancer | 10X | TPM | 120,300 | Clustering | Source | 200 | 20 | 5000 |
|  | Henry *et al.**^9^* | Human normal prostate | 10X | Raw count | 28,702 | Clustering | Target | 200 | 3 | 3000 |
| PBMC | Zheng *et al.**^10^* | Human PBMC | 10X | Raw count | 68, 579 | Clustering | Source | 200 | 20 | 5000 |
|  | Zheng *et al.**^10^* | Human PBMC | 10X | Raw count | 2,700 | Clustering | Target | 200 | 20 | 2000 |
| Pancreas | Baron *et al.**^11^* | Human pancreas | inDrop | Raw count | 8,562 | Clustering | Source | 200 | 20 | 2000 |
|  | Muraro *et al.**^12^* | Human pancreas | CEL-Seq2 | Raw count | 2,042 | Clustering | Source |  |  |  |
|  | Wang *et al.**^13^* | Human pancreas | SMARTer | Raw count | 430 | Clustering | Source |  |  |  |
|  | Segerstolpe *et al.**^14^* | Human pancreas | Smart-Seq2 | Raw count | 2,068 | Clustering | Target | 200 | 3 | 2000 |
| GBM | Neftel *et al.**^15^* | Human glioblastoma | 10X | Raw count | 12,074 | Algebra | Source | 200 | 20 | 1000 |
|  | Couturier *et al.**^16^* | Human glioblastoma | 10X | Raw count | 4,508 | Algebra | Target | 200 | 3 | 1000 |
|  | TCGA-GBM^17^* | Human glioblastoma | Bulk RNA-seq | Raw count | 613 | Algebra | Survival |  |  |  |
| BRCA | Wu *et al.**^18^* | Human breast cancer | 10X | Raw count | 24,271 | Algebra | Source | 200 | 20 | 1000 |
|  | Zhou *et al.**^19^* | Human Breast cancer | Smart-seq2 | Raw count | 545 | Algebra | Target | 200 | 3 | 1000 |
|  | TCGA-BRCA^17^* | Human Breast cancer | Bulk RNA-seq | Raw count | 1,079 | Algebra | Survival |  |  |  |
| mCRPC | He *et al.**^20^* | Human metastatic  prostate cancer | Smart-Seq2 | TPM | 836 | Algebra | Target | 200 | 3 | 1000 |
|  | Abida *et al.**^21^* | Human metastatic  prostate cancer | Bulk RNA-seq | TPM | 81 | Algebra | Survival |  |  |  |

Data downloaded from <https://www.cancer.gov/tcga>.

**Supplementary Table 3**: Summary of evaluation metric for all target datasets at resolution=1.0 analyzed in this paper

|  | **Normal Prostate** | | | **3K PBMC** | | | **Pancreas** | | | **In-house CITE-seq** | | |
| --- | --- | --- | --- | --- | --- | --- | --- | --- | --- | --- | --- | --- |
|  | **ARI** | **NMI** | **SI**  **(Cell type)** | **ARI** | **NMI** | **SI**  **(Cell type)** | **ARI** | **NMI** | **SI**  **(Cell type)** | **ARI** | **NMI** | **SI**  **(Cell type)** |
| **SCellBOW** | 0.26 | 0.56 | 0.13 | 0.49 | 0.52 | 0.11 | 0.56 | 0.82 | 0.53 | 0.65 | 0.72 | 0.06 |
| **Scanpy** | 0.16 | 0.53 | 0.13 | 0.36 | 0.5 | 0.1 | 0.38 | 0.73 | 0.24 | 0.51 | 0.66 | 0.08 |
| **Seurat** | 0.15 | 0.52 | 0.21 | 0.31 | 0.48 | 0.1 | 0.52 | 0.79 | 0.53 | 0.47 | 0.66 | 0.04 |
| **ItClust** | 0.01 | 0.02 | -0.05 | 0.33 | 0.46 | -0.05 | 0.31 | 0.29 | -0.05 | 0.43 | 0.55 | -0.13 |
| **scETM** | 0.11 | 0.37 | 0.04 | 0.38 | 0.5 | 0.08 | 0.35 | 0.67 | 0.46 | 0.41 | 0.61 | -0.03 |
| **scBERT** | 0.18 | 0.51 | 0.22 | 0.32 | 0.47 | 0.07 | 0.35 | 0.72 | 0.35 | 0.58 | 0.70 | 0.06 |
| **scPhere** | 0.04 | 0.42 | 0.05 | 0.19 | 0.38 | 0.04 | 0.23 | 0.63 | 0.33 | 0.24 | 0.55 | 0.21 |
| **DESC** | 0.17 | 0.52 | 0.02 | 0.46 | 0.49 | -0.06 | 0.39 | 0.72 | 0.2 | 0.54 | 0.65 | 0.01 |

**Supplementary Table 4:** Marker gene set major immune cell types

| **Major cell types** | **Marker genes** |
| --- | --- |
| B cells | *CD19, CD79A, MS4A1, CD74, HLA-DRA* |
| CD4 T | *IL7R, CCR7, CD3D, CD4* |
| CD8 T | *GZMK, CD8A, CD8B, GZMB* |
| DC | *CST3, CD14, ITGAM, ITGAX* |
| MAIT | *CD3D, KLRB1, RORA, ZBTB16* |
| Mono | *CD14, S100A12* |
| NK cells | *NKG7, GNLY, CD247, CCL3, GZMB, CD3D* |

**Supplementary Table 5:** Computation time across different transfer learning methods under the same hardware conditions (128 GB RAM, 16 core processor)

|  | **Wall time (Pancreas dataset)** | |
| --- | --- | --- |
|  | **Source Model** | **Target Model** |
| **SCellBOW (n_thread =1)** | 6min 21s | 2min 8s |
| **SCellBOW (n_thread =16)** | 2min 5s | 1min 20s |
| **ItClust** | 2min 4s | |
| **scETM (600 epoch, n_thread =16)** | 22min 38s | 5min 46s |
| **scETM (600 epoch, n_thread =1)** | 23min 49s | 5min 58s |
| **scBERT** | 3hrs 33min | 2min |

Supplementary Methods

#### **1. Datasets overview**

##### 1.1 Normal prostate data

We built our pre-trained normal prostate tissue model using the large Karthaus *et al*.^8^ dataset with approximately 120,300 cells as the source data. Cells in this study were isolated from histologically normal prostate regions of men treated for prostate cancer by radical prostatectomy. We obtained the processed TPM expression data from <https://singlecell.broadinstitute.org/single_cell/study/SCP864>, which was then log1p transformed, and 5,000 HVGs were selected for model training. We analyzed the clustering ability of the pre-trained model on the Henry *et al.**^9^* target dataset with approximately 28,702 cells from normal prostate specimens (young adult human prostate and prostatic urethra). The cells had been pre-annotated based on their similarity with matched single-cell transcriptomes of 9 FACS-purified cell subpopulations (Smooth Muscle (SM), Neuroendocrine epithelial (NE), Leukocytes (Leu), Luminal epithelial (LE), Hillock, Fibroblast (Fib), Endothelial (Endo), Club, and Basal epithelial (BE)). We retrieved the raw count data and cell type annotation from the GUDMAP database (<https://doi.org/10.25548/W-R8CM>). Raw counts were preprocessed according to ‘*Data preparation for clustering*’ in the main methods section, and 3,000 HVGs were selected for retraining. The 10x Chromium protocol was used for sequencing both source and target sample sets.

##### 1.2 Peripheral blood mononuclear cell (PBMC) data

We used a well-characterized reference 68K PBMC dataset from Zheng *et al*.^10^, with approximately 68,579 cells obtained fresh from a healthy donor as the source dataset. We used the 3K PBMC dataset from the same donor, consisting of 2,700 cryopreserved cells, as our target data. Both datasets are hosted on the 10x genomics website (<https://www.10xgenomics.com/>). We downloaded the gene-cell raw count matrix for the “Fresh 68k PBMCs (Donor A)” and “Frozen PBMCs (Donor A)” as our source and target datasets, respectively. We annotated the cell types in the target dataset based on similarity with expression profiles of 11 purified subpopulations of PBMCs (CD14+ Monocyte, CD19+ B, CD34+, CD4+ Memory T, CD4+ Naive T, CD4+ T Helper2, CD4+ T Reg, CD56+ NK, CD8+ Cytotoxic T, CD8+ Naive T, Dendritic) as a reference, as described by Zheng *et al*. Raw counts of both data sets were preprocessed according to ‘*Data preparation for clustering*’ in the main methods section. We selected a subset of 5,000 HVGs for the 68K PBMC and 2000 HVGs for the 3K PBMC dataset for further analysis.

##### 1.3 Pancreatic islet data

We used four single-cell RNA-seq (scRNA-seq) datasets (Baron *et al.**^11^*, Muraro *et al.**^12^*, Wang *et al.**^13^*, and Segerstolpe *et al.**^14^*) from the human pancreatic islet. The datasets are from multiple individual samples sequenced using different sequencing protocols. In the Baron dataset, the cells were sequenced using the inDrop protocol and are available on GEO: [GSE84133](https://www.ncbi.nlm.nih.gov/geo/query/acc.cgi?acc=GSE84133). The Muraro dataset is available with series number GEO: [GSE85241](https://www.ncbi.nlm.nih.gov/geo/query/acc.cgi?acc=GSE85241) and was sequenced using the CEL-Seq2 protocol. The Wang dataset is sequenced by the SMARTer protocol and is available under accession GEO: [GSE83139](https://www.ncbi.nlm.nih.gov/geo/query/acc.cgi?acc=GSE83139). The Segerstolpe dataset was sequenced by the Smart-Seq2 protocol and was retrieved from ArrayExpress (EBI) with accession number E-MTAB-5061. We used Baron, Muraro, and Wang datasets as our source data for the pre-trained model. A total of 11,181 cells were selected. We used the Segerstolpe dataset with 2,068 cells as the target dataset. Raw counts of both data sets were preprocessed according to ‘*Data preparation for clustering*’ in the main methods section, and 2000 HVGs were selected for further analysis. For these datasets, we used the cell labels given by the authors as ground truth. We filtered the unclassified and unclear cells from these datasets.

##### 1.4 Glioblastoma data

Glioblastoma multiforme (GBM) is the most aggressive, invasive, and undifferentiated type of tumor. Glioblastomas are highly heterogeneous and have well-characterized aggressive malignant subtypes: proneural (PN), classical (CL), and mesenchymal (MES)^22^. We have applied *phenotype algebra* to the GBM dataset from Couturier *et al.**^16^*, which consists of 4,508 malignant cells from one GBM patient (BT400). The 10X Genomics dataset sample BT400 was downloaded from <https://github.com/mbourgey/scRNA_GBM>. We clustered the scRNA-seq dataset using the Leiden clustering algorithm^23^ at a resolution of 1.0. We used well-known markers of cell types to perform cluster-level annotation of the dataset based on Gene Set Variation Analysis (GSVA)^24^ scores (see **Supplementary Methods** section 2.1). We have used the single-cell GBM dataset from Neftel *et al.**^15^* with 12,074 cells as our source data for transfer learning. The Neftel *et al.* 10X dataset was retrieved from GEO: [GSE131928](https://www.ncbi.nlm.nih.gov/geo/query/acc.cgi?acc=GSE131928). The source dataset was log normalized using Scanpy, and 1,000 HVGs were selected. For survival model training, we retrieved the survival and bulk RNA-seq expression data of the TCGA GBM subset with 613 samples from [https://portal.gdc.cancer.gov](https://portal.gdc.cancer.gov/). We used raw count expression matrices for all the datasets.

##### 1.5 Breast cancer data

We executed *phenotype algebra* on PAM50-subtyped breast cancer cells from the Zhou *et al.**^19^* study, with the raw count data available at GEO: [GSE118390](https://www.ncbi.nlm.nih.gov/geo/query/acc.cgi?acc=GSE118390). The authors annotated 545 malignant cells of six triple- negative breast cancer (TNBC) patients from Karaayvaz *et al.**^25^*, using the SubPred_pam50() function of the genefu R package^26^. As our source data for transfer learning, we used the single-cell TNBC dataset from Wu *et al.**^18^* with 24,271 cells from five patients, with the raw count data matrix downloaded from [https://singlecell.broadinstitute.org/single_cell/study/SCP1106/](https://singlecell.broadinstitute.org/single_cell/study/SCP1106/stromal-cell-diversity-associated-with-immune-evasion-in-human-triple-negative-breast-cancer). Raw counts were preprocessed according to ‘*Data preparation for clustering*’ in the main methods section, and 1000 HVGs were chosen for further analysis. For training the *phenotype algebra* survival model, we used the survival and bulk RNA-seq expression data of the TCGA BRCA subset from 1,079 samples retrieved from [https://portal.gdc.cancer.gov](https://portal.gdc.cancer.gov/).

##### 1.6 Metastatic prostate cancer data

We have used a publicly available metastatic castration-resistant prostate cancer (mCRPC) scRNA-seq data set originating from multiple donors and three metastatic tumor sites (lymph node, bone, and liver)^20^. In this dataset, we removed non-malignant cells based on the given cell type annotation published by the authors and focused further analysis on the 836 malignant cells derived from 11 patients. In prostate cancer, molecular subtypes are still poorly defined, and characterizing novel or more refined subtypes with clinical impact on patient prognosis is an active field of research. We employed GSVA scoring using established gene sets describing the molecular traits to classify 836 malignant cells. We annotated the tumor cells into one of the three categories: ARAH, NE, and ARAL (see **Supplementary Methods** section 2.2). We used the gene expression data and metadata from the publication. We retrieved survival and bulk RNA-seq expression data for 81 mCRPC patients provided by Abida *et al.**^21^* for training the survival model. We retrieved the dataset from [www.cbioportal.org](http://www.cbioportal.org) (prad_su2c_2019). We have used the Karthaus *et al*. pre-trained model as our source model for transfer learning. All samples available were individually normalized with transcripts per million (TPM).

#### **2. Molecular characterization of the cancer datasets**

##### 2.1 Annotation of Couturier *et al.* GBM dataset

For molecular characterization of the Couturier *et al.* datasets, we performed gene set enrichment analysis using GSVA scoring with default settings on the raw expression data. The gene sets comprise known marker genes^27^ for the Proneural (PRO), Classical (CLA), and Mesenchymal (MES) subtypes, as outlined in **Supplementary Table 6**.

**Supplementary Table 6:** Gene set for molecular subtypes of Glioblastoma

| **Subtype** | **Marker genes** |
| --- | --- |
| Proneural | *DLL3, BCAN, OLIG2, NCAM1, NKX2-2, ASCL1, PDGFRA* |
| Classical | *EGFR, CDKN2A, RB1, CDK4, CCDN2* |
| Mesenchymal | *CHI3L1, CD44, VIM, RELB, TRADD, PDPN, YKL40, MET, NF1, TNFRSF1A* |

We applied SCellBOW clustering on the dataset at a resolution of 1.0 and obtained 10 clusters. We performed a cluster-level subtype annotation based on the GSVA scores of the cells. For each cluster, we have assigned a molecular subtype based on the prevalent subtype (percentage of the cells) in that cluster. Clusters CL3, CL4, CL5, and CL6 have a higher percentage of PRO than MES and CLA and have thus been annotated as PRO. Similarly, clusters CL2, CL7, and CL8 were assigned MES, and clusters CL0, CL1, and CL9 were assigned CLA.

##### 2.2 Annotation of He *et al*. mCRPC dataset

For phenotypic characterization of the metastatic cells in the mCRPC target data, we performed GSVA scoring with default parameters on the preprocessed log1p(TPM) expression data. We used published marker gene sets (**Supplementary Table 7**) for ‘androgen receptor-positive prostate cancer’ (ARPC) and ‘neuroendocrine prostate cancer’ (NEPC) from Labrecque *et al.**^28^*, two well-described types of advanced prostate cancer.

**Supplementary Table 7:** Labrecque *et al*. 2019 gene sets for molecular subtypes of mCRPC

| **Subtype** | **Marker genes** |
| --- | --- |
| NEPC | *CHGA, SYP, ACTL6B, SNAP25, INSM1, ASCL1, CHRNB2, SRRM4* |
| ARPC | *AR, NKX3-1, KLK3, CHRNA2, SLC45A3, NAP1L2, S100A14, TRGC1, TARP* |

To obtain a high-level subgrouping, we performed K-means^29^ clustering on the ARPC and NEPC signature scores from GSVA using K-means++ initialization (**Supplementary Fig. 8a-b**). Based on the elbow method, the optimal number of clusters was determined to be 3. For further molecular characterization, we performed differential gene expression analysis on the obtained clusters using the *rank_genes_groups* function in Scanpy python library and, based on the results, annotated three subgroups of (i) ‘Androgen Receptor Activity High’ (ARAH), (ii) ‘Androgen Receptor Activity Low’ (ARAL), and (iii) ‘neuroendocrine-like’ (NE) cells. (**Supplementary Data 1**). The ARAH cluster was characterized by high expression of AR-activated (*TRGC1*, *TRGC2*, *KLK3*, *KLK2*, *GNMT*, *NKX3-1*, *NAP1L2*, *TMPRSS2*, *SLC45A3*, *C1orf116*) and NE-repressed genes (AR, SLC25A37, RGS10); while showing low expression of NE-activated genes (*SRRM4*, *SYP*, *SYT11*, *GNA01*, *KCNB2*, *GPX2*, *ETV5*, *INSM1*, *DNMT1*, *TRIM9*, *SNAP25*). The NE cluster exhibited high expression of NE-activated genes (*CHGA*, *SCG3*, *PROX1*, *ASCL1*, *SNAP25*, *TRIM9*, *INSM1*, *GPX2*, *EZH2*, *DNMT1*, *KCNB2*, *SYP*, *SYT11*, *SRRM4*); as wells as low expression of NE-repressed (*GATA2*, *RGS10*, *SLC25A37*, *MAPKAPK3*, *RIPK2*, *RGS10*) and AR activated genes (*ABCC4*, *CENPN*, *SLC45A3*, *NAP1L2*, *C1orf116*, *GNMT*, *FKBP5*, *PMEPA1*, *TRGC1*). The ARAL cluster had attenuated AR expression with measurable but low expression of some AR-activated (*GNMT*, *NAP1L2*, *TRGC1*, *TRGC2*) and NE-activated genes (*SYP*, *SYT11*, *GNAO1*, *GPX2*, *KCNB2*, *INSM1*, *PROX1*, *SCG3*, *ASCL1*, *CHGA*). Executing *phenotype algebra* on these broad subgroups resulted in a risk prediction matching the expectation and current knowledge.

#### **3. Gene set enrichment analysis on the mCRPC He *et al.* dataset**

For further characterization of the metastatic data set, not biased by previous knowledge or expectations, we performed SCellBOW clustering on the preprocessed in Scanpy log1p function on TPM expression data of the 836 malignant cells using resolution 0.8 and obtained eight clusters. We GSVA-scored as above, but against an extensive collection of ~16,500 gene sets describing a wide variety of cellular processes. We utilized gene sets from the Molecular Signatures Database (MSigDB.v.7.1:<http://www.gsea-msigdb.org/gsea/downloads_archive.jsp> H: hallmark gene sets, C2: curated gene sets and C5: ontology gene sets) in combination with in-house curated custom gene signatures (**Supplementary Data 1**). Performing differential gene set analysis across the SCellBOW clusters using rank_genes_groups function in Scanpy python library. with default settings using a Wilcoxon Rank Sum test and Bonferroni p-value adjustment) revealed activated and repressed pathways that were employed to describe the biology of clusters.

**Supplementary Note 1**

Comparison of SCellBOW clustering efficacy with additional methods

For benchmarking, we further compared the performance of SCellBOW with two recently published tools- scPhere^6^ and scBERT^5^. scBERT handles scRNA-seq data by leveraging contextualized embeddings built upon BERT^30^, which capture the context and relationships between genes in a cell's transcriptional profile. Its primary emphasis lies in cell annotation, particularly focusing on cell type classification. On the other hand, scPhere is a deep generative model that embeds cells into geometric spaces, enabling a unique way of representing cells within specialized geometric dimensions compared to traditional Euclidean spaces. For comparing the clustering ability, we used the Normal prostate, pancreatic islet, 3K PBMC, and our in-house PBMC-spleen datasets for our benchmarking (see **SCellBOW robustly dissects tissue heterogeneity** in the main text).

We computed the adjusted Rand index (ARI) for a resolution ranging from 0.2 to 2.0 in 0.2 intervals and found that SCellBOW consistently outperformed scBERT and scPhere across all resolutions (**Supplementary Fig. 3a-d**). Then, for the same range of resolutions, we computed the normalized mutual information (NMI) (**Supplementary Fig. 3e**) and observed that scPhere consistently exhibited poorer performance compared to SCellBOW and scBERT across all datasets. SCellBOW outperformed scBERT across all public datasets. In terms of the Silhouette Index (SI) for cell types, SCellBOW demonstrated superior performance over scPhere and scBERT in the 3K PBMC and Pancreas datasets (**Supplementary Fig. 3f**). However, in the in-house CITE-seq dataset, scPhere displayed improved results, while scBERT exhibited enhanced performance in the normal prostate dataset.

#### **Supplementary Note 2**

##### Processing of in-house PBMC-spleen dataset in SCellBOW

Dataset: This CITE-seq dataset includes 4,819 cells and 33,538 genes from the spleen and matching PBMC of 4 polytraumatized patients with splenectomy hemostasis, generated by us using 10X.

Cell and gene filtering criteria: 1) eliminated cells with gene counts <200 using filter_cells() function from Scanpy Python package; 2) eliminated genes if the number of cells expressing this gene is < 3 filter_genes() function.

Data processing: 1) gene expression levels for each cell were normalized using the normalize_total() function in Scanpy with target_sum=10,000; 2) normalized gene expression was then transformed using log(1+x) transformation with natural logarithm using log1p function; 3) top 3,000 highly variable genes were selected using the highly_variable_genes() function in Scanpy; 4) the expression is further standardized to a z-score using scale function, and the standardized gene expression values were used as input for SCellBOW. After the above filtering and data processing, 4,785 cells and 3,000 highly variable genes remained for downstream analysis.

Clustering and visualization using SCellBOW: All 4,785 cells were clustered by SCellBOW with default parameters using the leiden() function. We used the umap to visualize the 2D plot of the clusters in SCellBOW.

Cell type labeling of CITE-seq using Azimuth: Cell type annotation was performed using Seurat-based Azimuth^31^. The molecular reference atlas for Human-PBMC was used for annotation from <https://azimuth.hubmapconsortium.org/>. We eliminated the cells with a cell type count< 5. There were 4,817 remaining cells.

**Supplementary Note 3**

##### Erythroid cells in the PBMC-spleen dataset

In our analysis, we observed that cells identified as ‘Eryth’ by Azimuth are scattered across all the clusters. Since erythroid cells were removed during the CITE-seq experiments, erythroid cells are not part of PBMCs in this dataset. Thus, we assumed that if erythrocytes are present, it is most likely due to contamination. Since erythroid cells form a majority of cluster CL6 amidst T cells, they are putative ambient RNA contaminations misguiding the Azimuth annotation (**Supplementary Fig. 4e**). We further performed differential gene expression analysis based on cell types using the Scanpy rank_genes_groups function on the PBMC-spleen dataset. We observed that they have mitochondrial genes as highly differentially expressed genes. This can be attributed to the process of erythropoiesis in cells^32^. However, given the spatial distribution of cells classified as ‘Eryth’ across multiple clusters combined with high mitochondrial gene content in the cells, we assume that ‘Eryth’ cells are misclassified/unclassified cells (**Supplementary Fig. 4g**).

#### **Supplementary Note 4**

##### Performance of *phenotype algebra* using fixed-length embedding from existing tools

##### To illustrate the effectiveness of fixed-length embeddings obtained from SCellBOW for *phenotype algebra,* we compared them to other tools such as scETM^31^, scPhere^6^, and scBERT^5^. We used the GBM, BRCA, and mCRPC datasets (see **Description of datasets for model evaluation** under Methods section) to compare the accurate relative ranking ability for our benchmarking (see **SCellBOW facilitates subclonal survival-risk attribution of tumors** in the main text). Unlike SCellBOW, which is an unsupervised transfer learning-based model, the limitation of scETM and scBERT is the dependency on the cell type labeling of the source dataset. Due to the lack of source data annotation in Neftel *et al.* GBM dataset, we used the Scanpy clusters as the source annotation. scPhere is a deep generative model that embeds cells into low-dimensional space and doesn’t need any source data. In the case of scPhere and scBERT, we have used embedding size = 300 to match SCellBOW. However, in the case of scETM, we adhered to the default embedding size of 50, since it does not explicitly allow the tuning of this parameter.

##### In the case of GBM and mCRPC, we observed that all the tools (scETM, scPhere, and scBERT) failed to detect the relative position of the subtypes in the combined state ((MES+CLA) for GBM and (ARAH+ARAL+NE) for mCRPC) (**Supplementary Fig. 6**). While scETM and scPhere correctly identified the least aggressive subtype in GBM as PRO; it failed to position CLA and MES accurately (**Supplementary Fig. 6a-c**). Similarly, in BRCA, we observed that scETM and scPhere could identify LUMA as the least aggressive subtype, however, they failed to detect the relative position of the remaining subtypes (**Supplementary Fig. 6d-f**). scBERT outcomes didn't align with the established aggressiveness order both in GBM and BRCA. In mCRPC, the known aggressiveness order is NE $\succ$ ARAL $\succ$ ARAH. scETM, scPhere, and scBERT disagreed with the known relationship among these subtypes (**Supplementary Fig. 6g-i**). scETM, scBERT, and scPhere struggle to differentiate cancer clones based on their aggressiveness and their impact on disease prognosis.

**Supplementary Note 5**

##### The importance of choosing Doc2vec as our base model

Single-cell RNA-seq profiles can relate cellular states and mRNA expression relationships, revealing gene expression programs corresponding to tumorigenesis. Identifying the groups of cells with similar phenotypic profiles is a critical step in cellular heterogeneity dissection. Natural language is a valuable analogy for cell signals where cells are analogous to sentences, and genes are analogous to words. Any downstream analysis of the scRNA-seq requires data to be represented in fewer dimensions. Embeddings are fixed-length vector representations. Using the cell sentences analogy, SCellBOW creates single-cell embedding based on the natural language processing (NLP) - Doc2vec language model^33^. We have tested the performance of SCellBOW compared to Scanpy and Doc2Vec^34^ on Segerstolpe *et al.* pancreas data. As source data for SCellBOW pre-trained model, we have taken Baron *et al.* Both datasets were retrieved from ItClust (<https://github.com/jianhuupenn/ItClust>).

In **Supplementary Fig. 7**, we observed that the majority of cells of similar cell types are more localized in the 2-dimensional space compared to other Scanpy and Doc2vec. After SCellBOW, Doc2vec was able to group cells with similar signatures closer to each other (**Supplementary Fig. 7g-i**). Thus, embedding generated by the Doc2vec-based approaches preserves spatial segregation across the clusters. We observed the highest overall quality scores (ARI, NMI, and cell type SI) in SCellBOW (see **Supplementary Fig. 7j**). The SCellBOW score being higher than the Doc2Vec embedding and Scanpy embeddings suggest that although the Doc2vec model has good performance compared to Scanpy, transfer learning using the Doc2vec model yields higher clustering accuracy.

#### **Supplementary Note 6**

##### Extended analysis of differentially expressed genes in spleen against PBMC

Spleen is the largest secondary lymphoid organ in the body and orchestrates a wide range of immunological functions. The physical organization of the spleen allows it to filter blood-borne pathogens and antigens. Differential expression analysis between immune cells originating from the spleen and peripheral blood exhibited spleen-specific elevated expression of dissociation-associated genes (*JUNB*, *FOSB*, *JUND*, *JUN*) and stress response heat shock genes (*HSP*, *HSP90AB1*, *HSPD1*, *DNAJB1*) (**Supplementary Fig. 4h,i and Supplementary Data 2**). Upregulation of dissociation and stress response-associated gene families can be attributed to the additional dissociation step during sample preparation, which is exclusive to the splenocytes^35,36^. Furthermore, B cell differentiation-related genes such as *VPREB3*, *MS4A1*, and *CD83* were found upregulated among splenocytes as compared to PBMC, suggesting the dominance of various stages of development, differentiation, and maturation of B cells in the spleen. Notably, the spleen has different structures, such as primary follicles and the germinal center within the white pulp where various stages of B cells reside. We also spotted spleen-specific upregulation of nuclear receptors (*NR4A1*, *NR4A2*) and B cell receptors (BCR) (*BANK1*, *CD79A*, *CD79B*), indicating that B cells are inherently different between circulation and the spleen and likely undergo additional maturation steps in the spleen. We also observed spleen-specific expression of B cell inhibitory receptors (BCIR) (*CD22*, *CD72*, *LY9*), suggesting that multipotent progenitors reside in the spleen. These multipotent progenitors of the spleen stem cells could be differentiating to myeloid and erythroid lineage cells in addition to differentiation of the lymphoid lineage cells (primarily to B cells and to some extent to T cells)^37^. Overall, the BCR and BCIR are two important receptors that play complementary roles in regulating B cell activation and tolerance. While the BCR initiates signaling cascades that lead to B cell activation, the BCIR limits these signals and helps maintain B cell tolerance.

B cells are capable of developing a wide range of effector functions, including antibody secretion via differentiating to plasma cells, antigen processing and presentation to T helper cells, cytokine production, and generation of immunological memory. The mature spleen plays an important role in B cell development and antigen-dependent maturation because it is the site of terminal differentiation for developing B cells that arrive from the bone marrow known as Transitional B cells of type 1 (T1), and these cells develop to transitional B cells of type 2 (T2) and reside to the primary follicles of the spleen^38^. Naive B cells enter the spleen from the circulation, where they differentiate into either B effectors that differentiate into antibody-secreting plasma cells or B memory cells^39^. The prominent role of the spleen in B cell differentiation and activation inspired us to delve deeper into compositional and phenotypic differences associated with B cell subtypes across peripheral blood and spleen. Based on annotation, as expected, we noted enrichment of B effectors in the spleen (**Supplementary Fig. 4j**). SCellBOW embedding could spatially segregate B effectors, B naive cells, and plasmablasts, although effectors and naive cells were not split into two distinct clusters (**Supplementary Fig. 4k**).

Further analysis of B cell subtypes showed higher expression of the dissociation genes and stress response of genes in spleen B effector and B naive cells as compared to their peripheral blood counterpart (**Supplementary Fig. 4l, m)**. B naive has higher expression of actin genes (*ACTB*, *ACTG1*) as compared to B effectors, suggesting the naive B cells are more dynamic for differentiation to any of B cell subtypes, such as the effector or memory^40^. Notably, the mutation in these genes results in the development of large B-cell lymphoma. In the case of the B effector, we observed that the expression of *CD27*, *HLADQA1*, *CD83*, *PRMT1*, and *CD69* are significantly higher in the mature effector B cells^41,42^. Collectively, genes upregulated in the B cells from the spleen were found to be involved in cell-cell adhesion, activation of the B cell receptor signaling pathway, and differentiation of B cells during hematopoiesis and our analysis very well corroborates with known B cell development biology under physiological conditions.

#### **Supplementary Note 7**

##### Dependency of *phenotype algebra* on SCellBOW embeddings

The concept of *phenotype algebra* is new, and there is no algorithm that is directly comparable to *phenotype algebra* on the specific task of sub-clonal survival risk (aggressiveness) stratification. However, *phenotype algebra* is built upon fixed-length embeddings generated in SCellBOW. To assess the dependency of *phenotype algebra* on these embeddings, we examined the possibility of achieving risk inference directly from gene expression data for known molecular subtypes of specific cancers (BRCA, GBM, mCRPC).

In the case of GBM, we observed that the gene expression model failed to correctly position CLA, MES, and (CLA+MES) in terms of their aggressiveness (**Supplementary Fig. 8i**). For BRCA, while it effectively positioned LUMA and HER2, it incorrectly categorized LUMB as the most aggressive subtype, despite LUMB typically having a favorable prognosis (**Supplementary Fig. 8j**). Similarly in mCRPC, the gene expression-based model identified the NE subtype as the least aggressive, contradicting established literature, which recognizes NE as the most aggressive subtype (**Supplementary Fig. 8k**). These results demonstrated that using the survival modeling techniques directly on gene expression data, as opposed to SCellBOW embeddings, failed to accomplish accurate risk stratification.

### Supplementary Figures

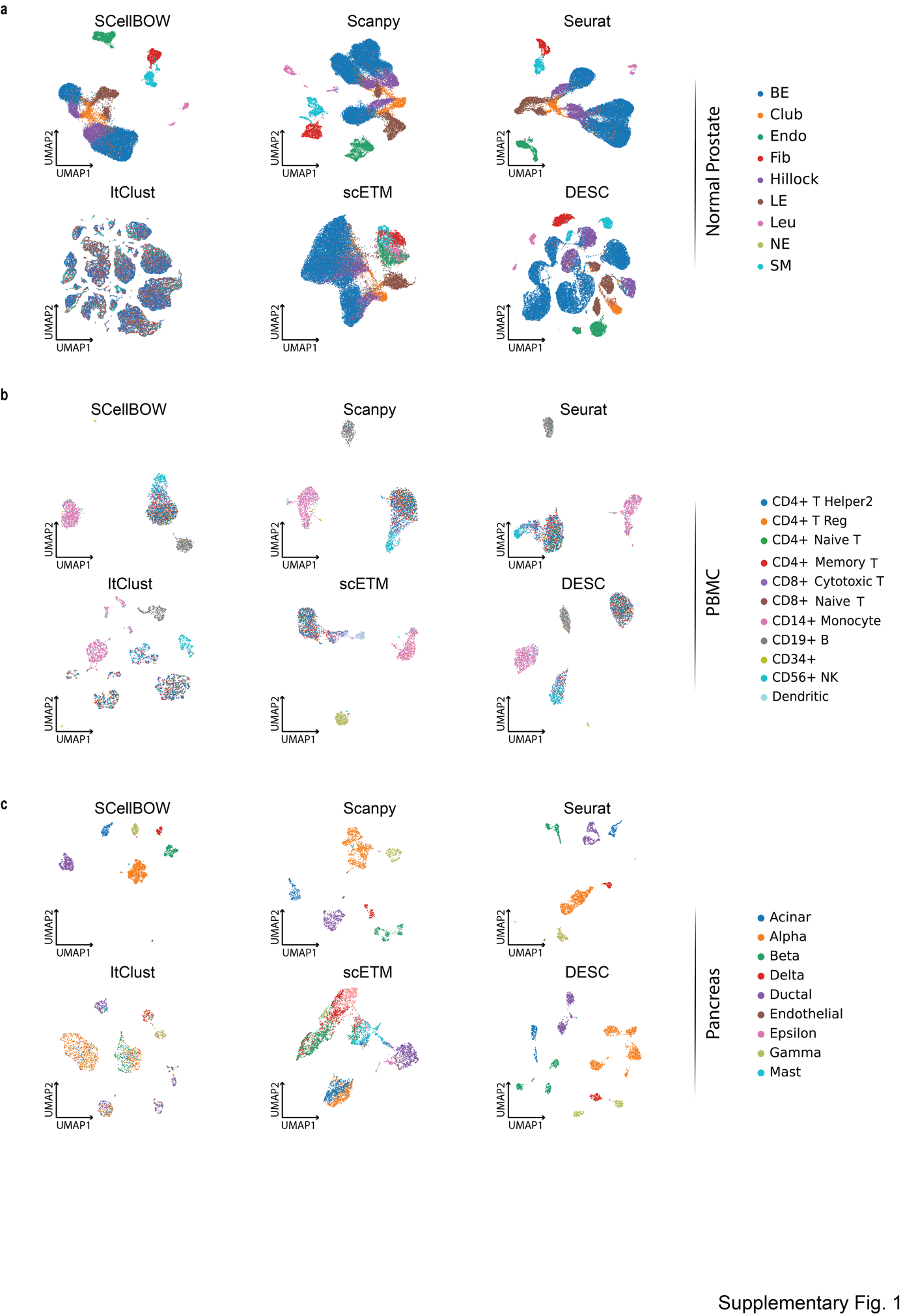

**Supplementary Fig. 1**. **Cell embeddings visualization**

**a-c**, The UMAP plots showing embedding of SCellBOW compared to different existing methods benchmarked on normal prostate (a), peripheral blood mononuclear cells (PBMC) (b), and pancreas datasets (c). The coordinates of all the plots are colored by true cell types.

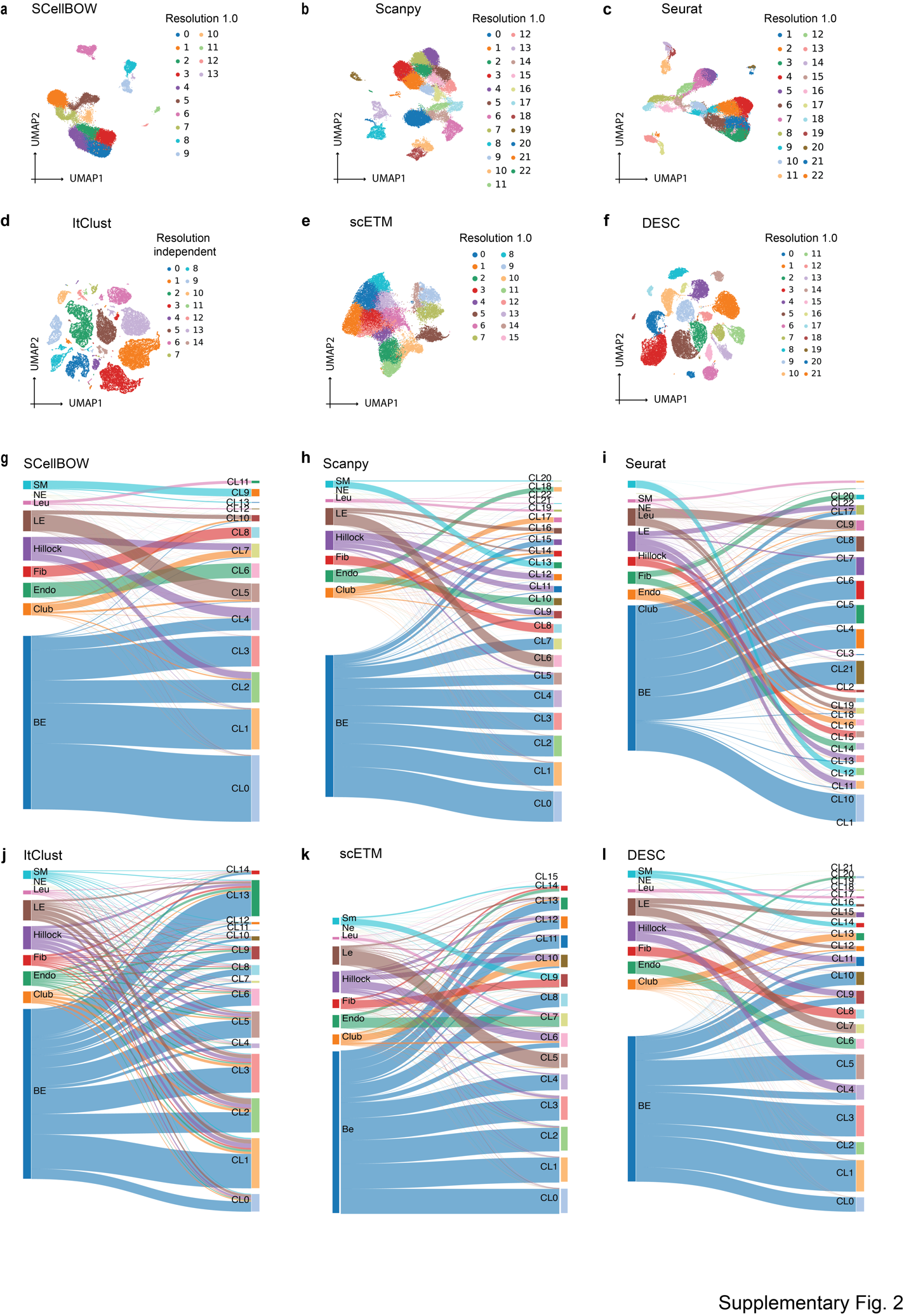

**Supplementary Fig. 2**. **Cell embedding visualization of the normal prostate scRNA-seq dataset.**

**a-f,** The UMAP plots showing embedding of SCellBOW, Scanpy, Seurat, ItClust, ProjectR, and DESC on normal prostate. The coordinates of all the plots are colored by clusters.

**g-l,** Alluvial plots showing the mapping of clusters resulting from the benchmarking tools onto the true cell types from Henry *et al*. normal prostate dataset. CL is used as an abbreviation for cluster.

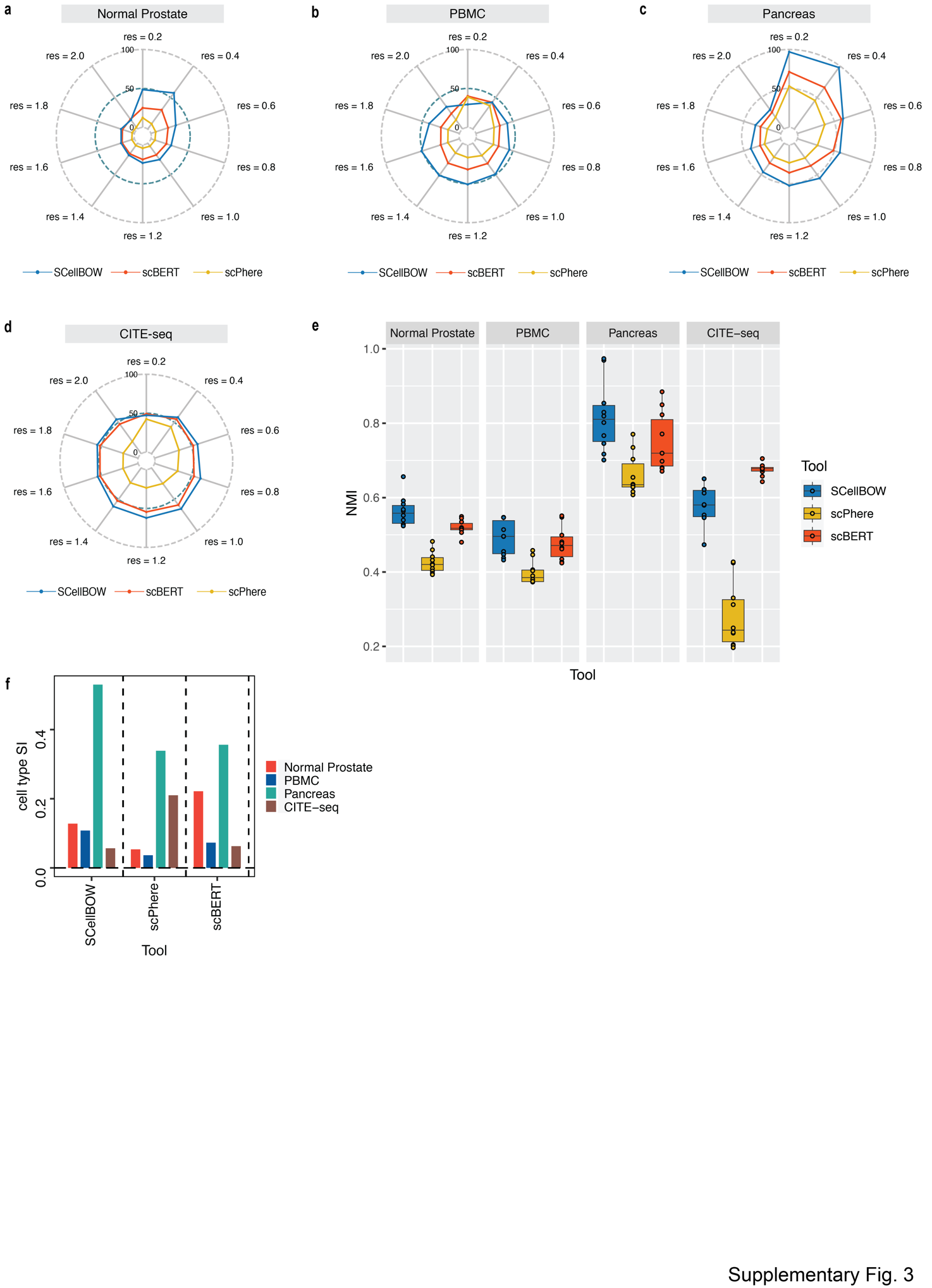

**Supplementary Fig. 3**. **Extended performance evaluation of SCellBOW with scBERT and scPhere.**

**a-d,** Radial plot for the percentage of contribution of different methods towards ARI for various resolutions ranging from 0.2 to 2.0 for normal prostate (a), PBMC (b), pancreas (c), and CITE-seq (d) datasets.

**e,** Box plot for the NMI of different methods across different resolutions ranging from 0.2 to 2.0 in steps of 0.2.

**f,** Bar plot for the cell type silhouette index (SI) for different methods. The default resolution was set to 1.0.

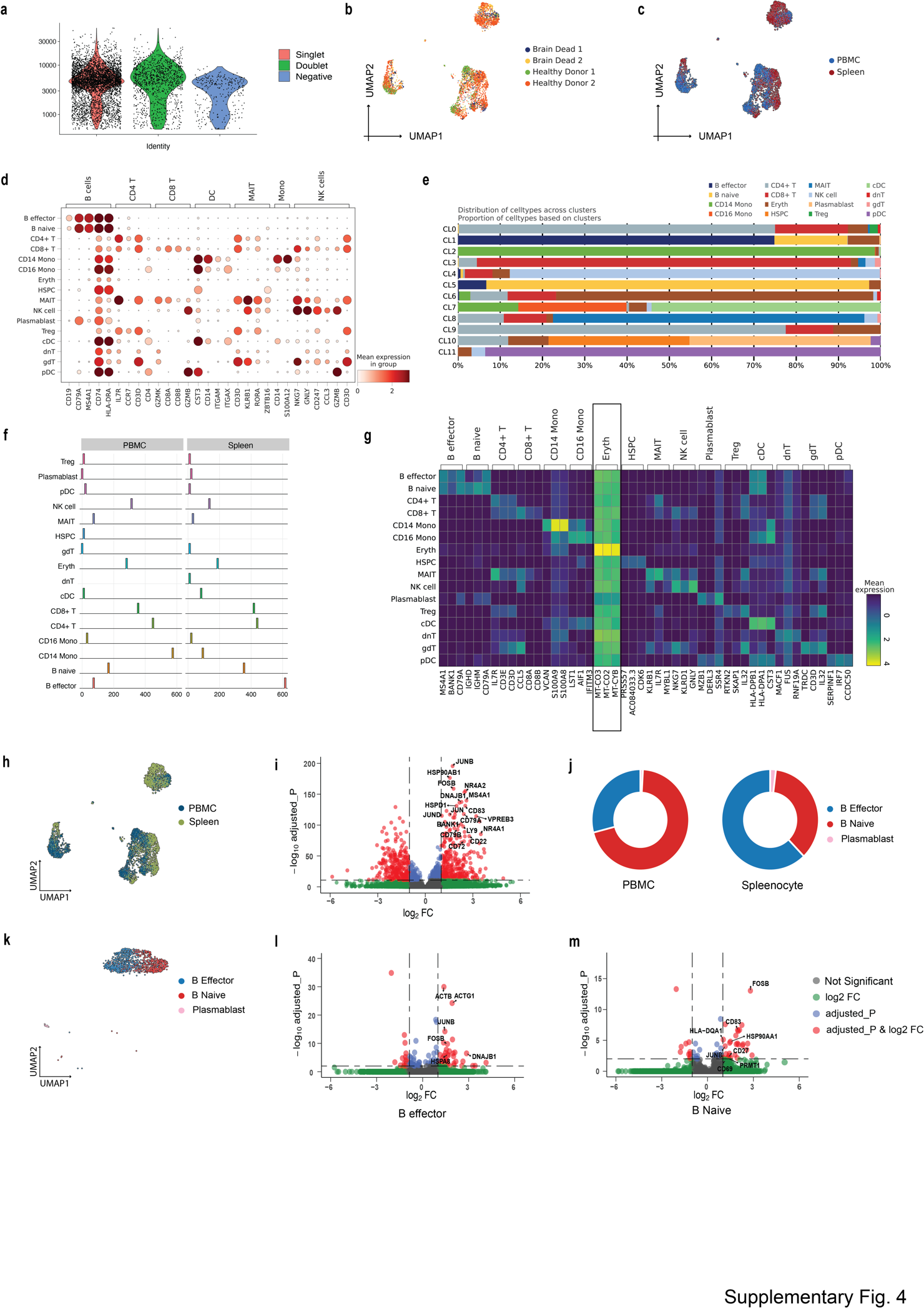

**Supplementary Fig. 4**. **Extended analysis of in-house CITE-seq dataset.**

**a,** Violin plot showing the distribution of UMIs for singlets, doublets, and negative cells.

**b,** The UMAP plots showing embedding of SCellBOW colored by donors.

**c,** The UMAP plots showing embedding of SCellBOW colored by tissue of origin.

**d,** Dot plot to check the expression of marker genes of PBMC per cell type identified by Azimuth.

**e,** Bar plot showing the proportion of annotated cell types across different clusters of SCellBOW.

**f,** Compositional difference in proportion annotated cell types in PBMC vs. splenocytes.

**g,** Heatmap for annotated cell type-wise differentially expressed genes in each cell type.

**h,** UMAP plots of the cells colored by their tissue source.

**i,** Volcano plot showing the differential genes (red dots) in the spleen and PBMC for B cells (P-value <0.05, False discovery rate (FDR) <0.01).

**j,** Donut plot showing the compositional difference in the proportion of B cell subtypes (B naive, B effector, and plasmablast) in PBMC and spleen.

**k,** UMAP plot showing the embedding of SCellBOW colored by B cell subtypes.

**l-m,** Volcano plot showing the differential genes (red dots) in the spleen and PBMC for B effector (l) and B naive cells (m) (P-value <0.05, FDR <0.01).

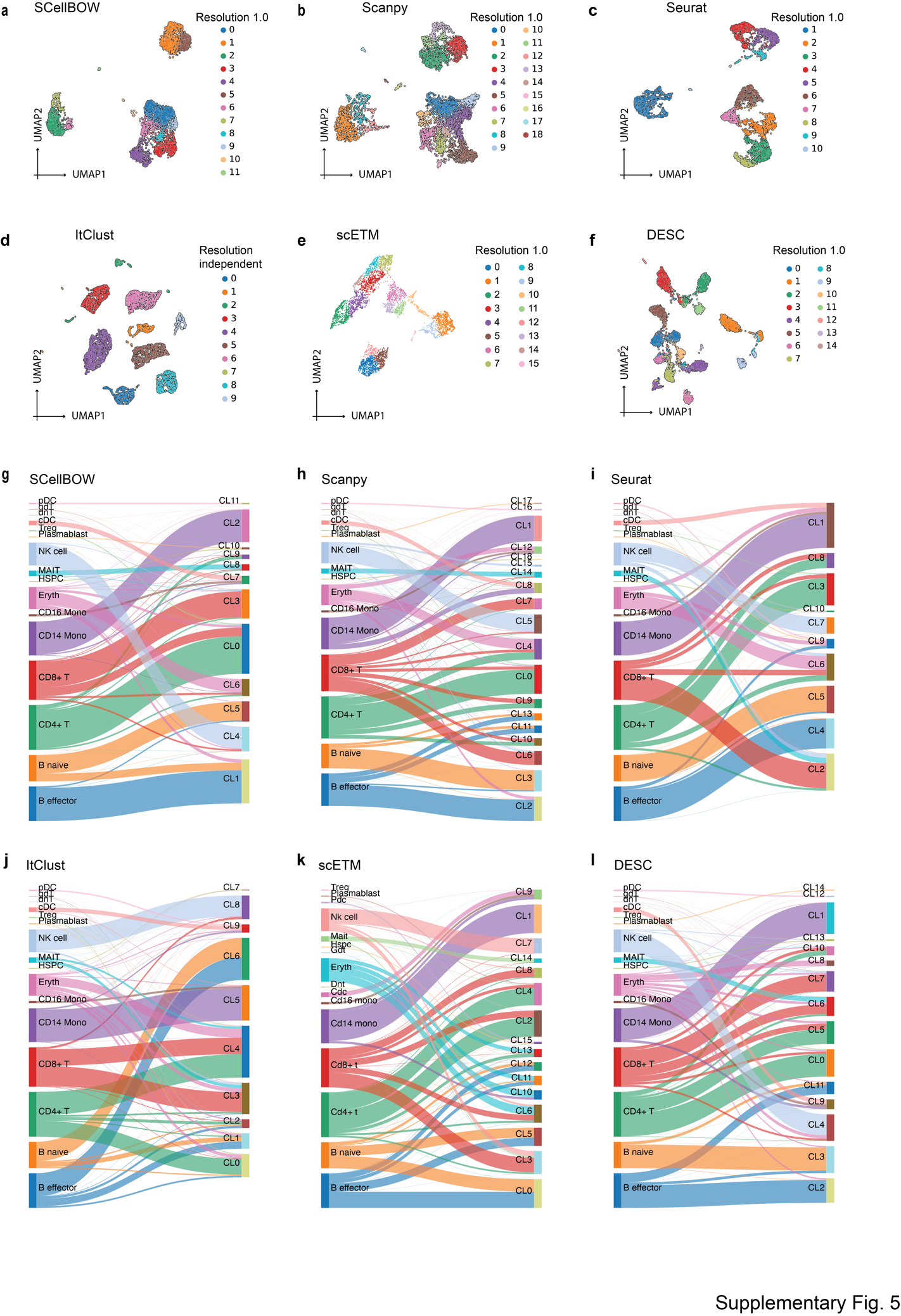

**Supplementary Fig. 5. Cell embedding visualization of the in-house CITE-seq scRNA-seq dataset.**

**a-f,** The UMAP plots showing embedding of SCellBOW, Scanpy, Seurat, ItClust, ProjectR, and DESC on the PBMC-spleen dataset. The coordinates of all the plots are colored by clusters.

**g-l,** The alluvial plots showing the mapping of clusters resulting from the benchmarking tools onto the cell types identified by Azimuth.

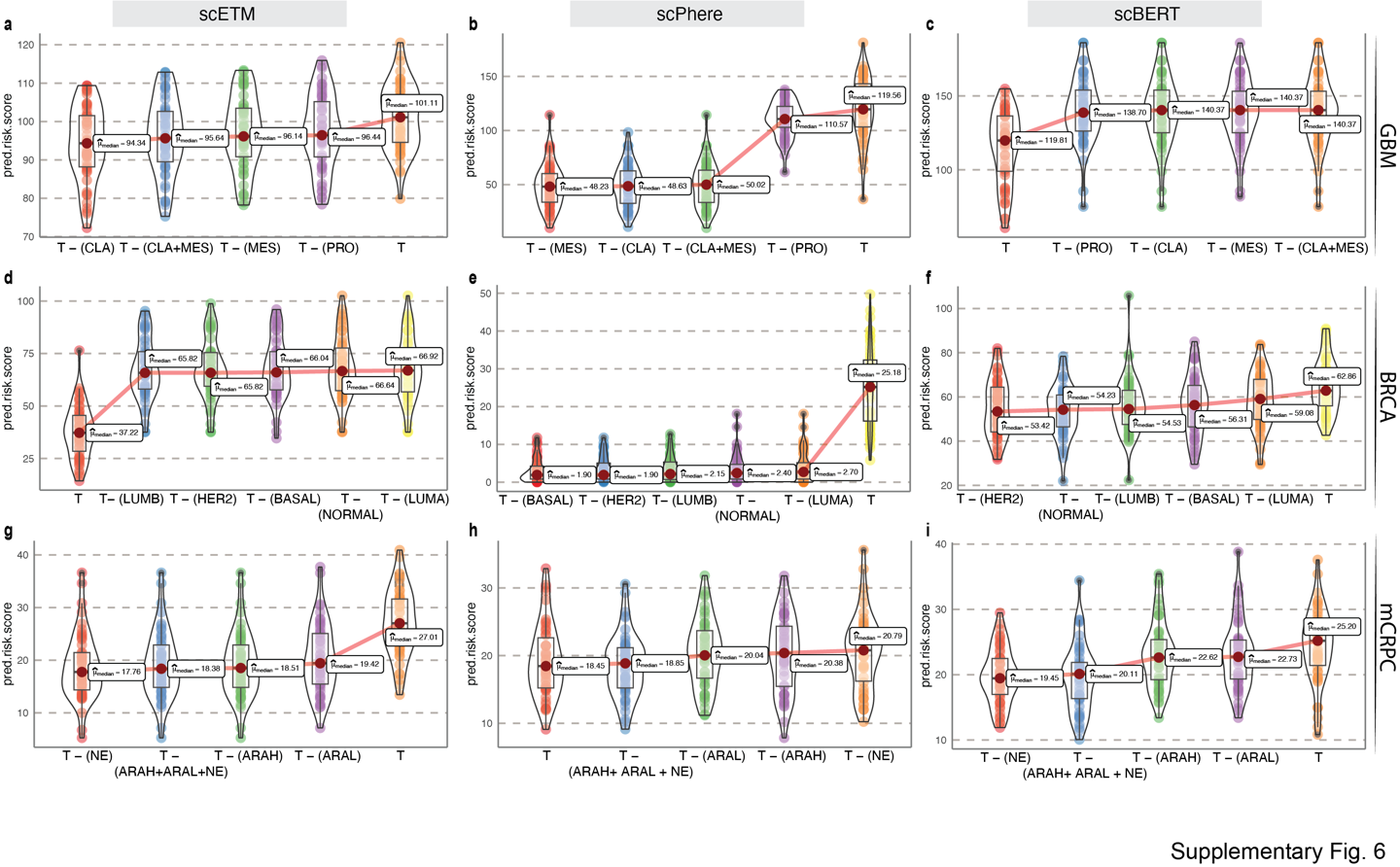

**Supplementary Fig. 6**. **Survival risk inference using fixed-length embeddings** **from scETM, scPhere, and scBERT.**

**a-c,** *Phenotype algebra*-based risk scores of GBM molecular subtypes using fixed-length embeddings from scETM (a), scPhere (b), and scBERT (c). The *total tumor* is denoted by *T*.

**d-f,** *Phenotype algebra*-based risk scores of PAM50 molecular subtypes of BRCA using fixed-length embeddings from scETM (a), scPhere (b), and scBERT (c).

**g-i,** *Phenotype algebra*-based risk scores of three high-level categories of mCRPC using fixed-length embeddings from scETM (a), scPhere (b), and scBERT (c).

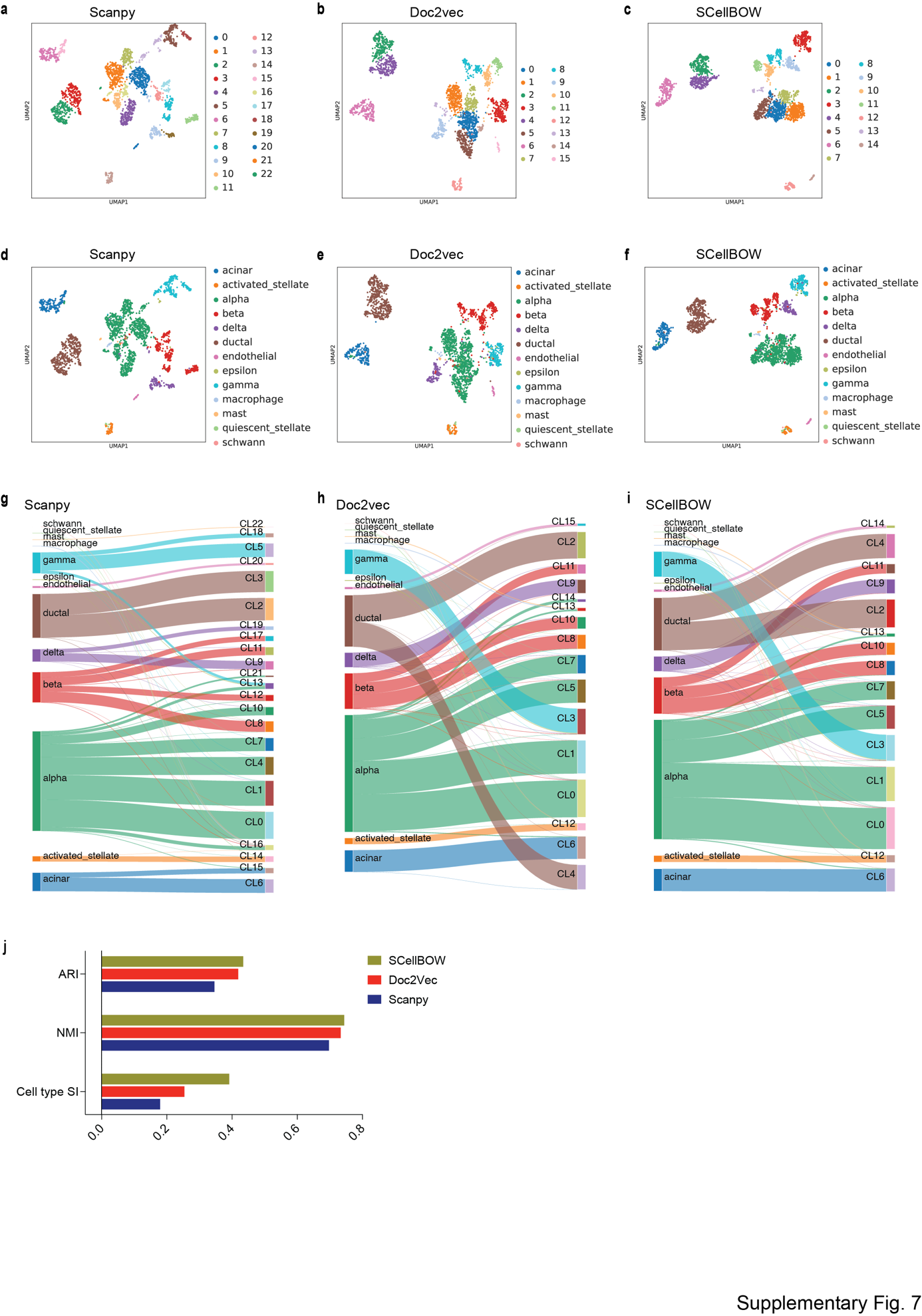

**Supplementary Fig. 7**. **Assessing the quality of clustering using transfer learning on BOW models using the pancreas dataset.**

**a-c,** UMAP plot of Scanpy embedding (a), Doc2Vec embedding (b), and SCellBOW embedding (c) of pancreas dataset using Leiden clustering at resolution 1.0.

**d-f,** UMAP plot of Scanpy embedding (d), Doc2Vec embedding (e), and SCellBOW embedding (f) of pancreas dataset colored with their annotated cell types

**g-i,** Alluvial plot for cell types against Leiden clusters for Scanpy (g) Doc2vec (h) SCellBOW (i).

**j,** Barplot for ARI, NMI, cluster purity, Silhouette index (cell type and cluster).

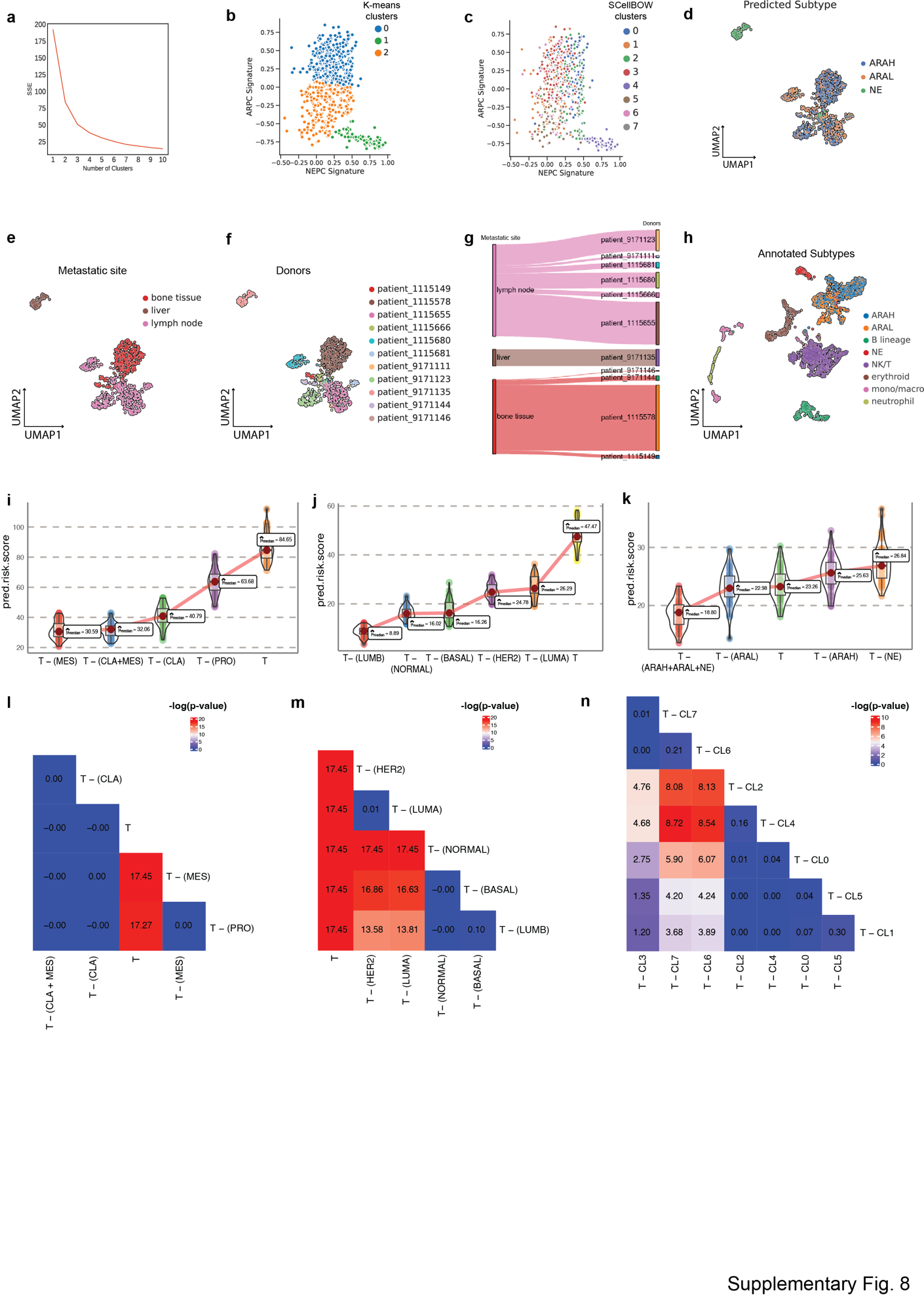

**Supplementary Fig. 8**. **Extended analysis of He *et al.* single-cell mCPRC dataset.**

**a,** Elbow plot for selecting the best K for K-means clustering.

**b-c,** Scatter plot of GSVA scores of ARPC and NEPC gene sets colored by the K-means clusters (E) and SCellBOW clusters (F).

**d,** UMAP plot visualizing the high-level ARAH, ARAL, and NEPC categories on SCellBOW embeddings.

**e-f,** The UMAP plots showing the embedding of SCellBOW colored by metastasis site (a) and donors (b).

**g,** Alluvial plot to visualize tumor metastasis site of the donors.

**h,** UMAP plot visualizing SCellBOW embeddings of tumor microenvironment cells (malignant + non-malignant) in the He et al. dataset based on author annotations.

**i-k,** *Phenotype* *algebra*-based risk scores using gene expression profile of GBM molecular subtypes (h), PAM50 molecular subtypes of BRCA (i), three high-level categories of mCRPC (j). The *total tumor* is denoted by *T*.

**l-n,** Heatmap for -log_10_(p-value) of the predicted risk scores for GBM subtype (k), BRCA subtype (l), and mCRPC clusters (m), and using Wilcoxon unpaired one-sided test.

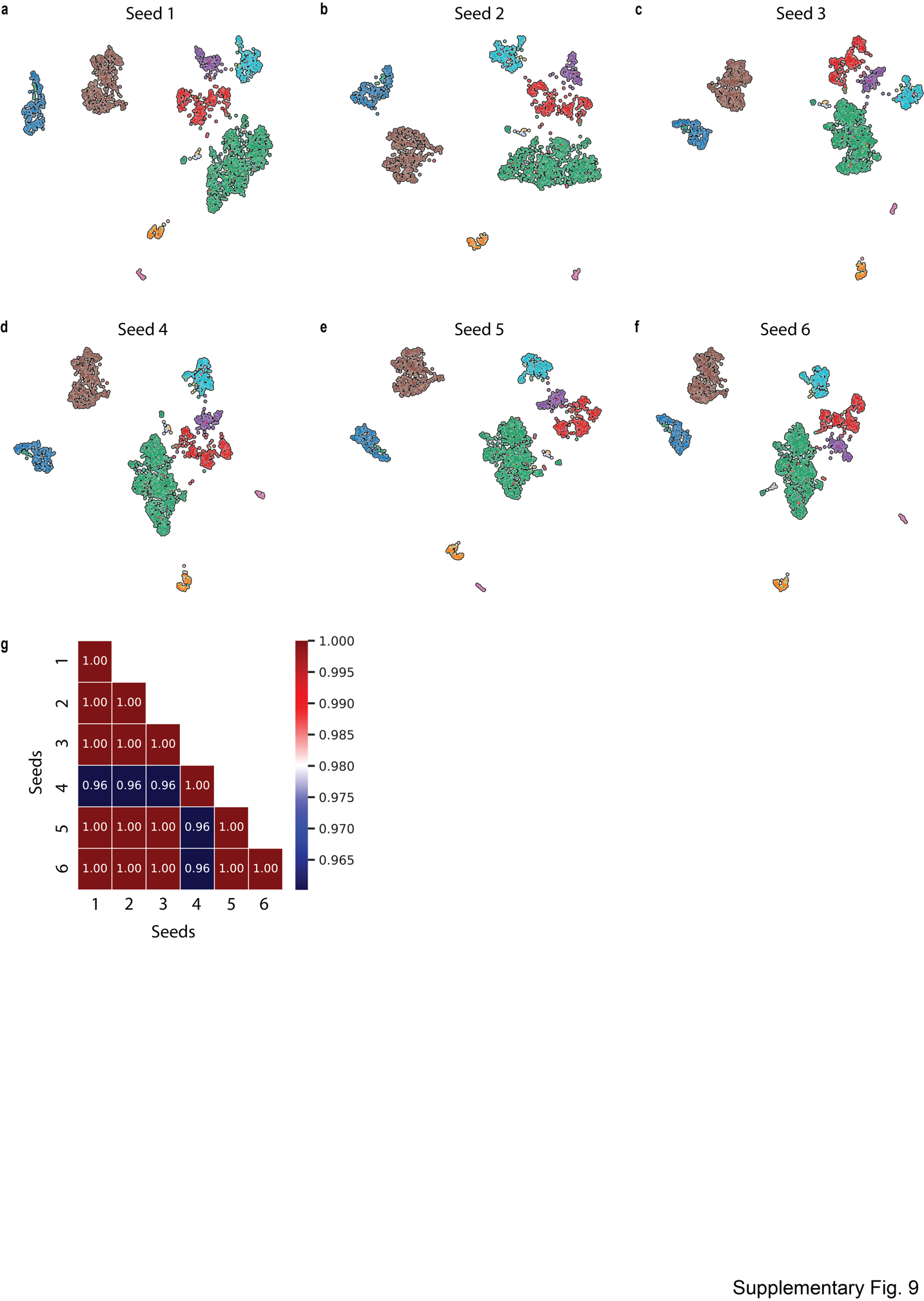

**Supplementary Fig. 9**. **Assessing the effect of random seed on the quality of cell sentence generation using the pancreas dataset.**

**a-f,** The UMAP plots showing the embedding of SCellBOW generated using cell sentences with different random seeds ranging from 2 to 25. The colors of the cells in the UMAP plots indicate clusters.

**g,** Heatmap showing the ARI between each pair of clustering outcomes with distinct seeds.
